# Effects of gene dosage on cognitive ability: A function-based association study across brain and non-brain processes

**DOI:** 10.1101/2024.04.16.589618

**Authors:** Guillaume Huguet, Thomas Renne, Cécile Poulain, Alma Dubuc, Kuldeep Kumar, Sayeh Kazem, Worrawat Engchuan, Omar Shanta, Elise Douard, Catherine Proulx, Martineau Jean-Louis, Zohra Saci, Josephine Mollon, Laura M Schultz, Emma E M Knowles, Simon R. Cox, David Porteous, Gail Davies, Paul Redmond, Sarah E. Harris, Gunter Schumann, Guillaume Dumas, Aurélie Labbe, Zdenka Pausova, Tomas Paus, Stephen W Scherer, Jonathan Sebat, Laura Almasy, David C Glahn, Sébastien Jacquemont

## Abstract

Genomic Copy Number Variants (CNVs) that increase risk for neurodevelopmental disorders are also associated with lower cognitive ability in general population cohorts. Studies have focussed on a small set of recurrent CNVs, but burden analyses suggested that the vast majority of CNVs affecting cognitive ability are too rare to reach variant-level association. As a result, the full range of gene-dosage-sensitive biological processes linked to cognitive ability remains unknown.

To investigate this issue, we identified all CNVs >50 kilobases in 258k individuals from 6 general population cohorts with assessments of general cognitive abilities. We performed a CNV-GWAS and functional burden analyses, which tested 6502 gene-sets defined by tissue and cell-type transcriptomics as well as gene ontology disrupted by all rare coding CNVs.

CNV-GWAS identified a novel duplication at 2q12.3 associated with higher performance in cognitive ability. Among the 864 gene-sets associated with cognitive ability, only 11% showed significant effects for both deletions and duplication. Accordingly, we systematically observed negative correlations between deletion and duplication effect sizes across all levels of biological observations. We quantified the preferential effects of deletions versus duplication using tagDS, a new normalized metric. Cognitive ability was preferentially affected by cortical, presynaptic, and negative-regulation gene-sets when duplicated. In contrast, preferential effects of deletions were observed for subcortical, post-synaptic, and positive-regulation gene-sets. A large proportion of gene-sets assigned to non-brain organs were associated with cognitive ability due to low tissue specificity genes, which were associated with higher sensitive to haploinsufficiency. Overall, most biological functions associated with cognitive ability are divided into those sensitive to either deletion or duplications.

## Introduction

Copy Number Variants (CNVs) are deletions or duplications larger than 1000 base pairs^1^. CNVs are major contributors to risk for neurodevelopmental disorders^2^, including intellectual disability (ID)^3–5^, autism spectrum disorder (ASD)^6–8^, and schizophrenia^8–10^. CNVs that increase the risk for psychiatric conditions also appear to invariably affect cognitive abilities and are often associated with multi-morbidity in the clinic^11–13^. The routine implementation of whole genome CNV detection, as a first-tier diagnostic test, identifies “pathogenic” CNVs in 10 to 15 % of children with neurodevelopmental disorders (NDD)^14^. But beyond the benign vs pathogenic categorical classification of genomic variants, their effect size on cognitive ability has been used to provide more nuanced information on the severity of a variant and to quantify the risk for NDD. Cognitive ability remains one of the traits most commonly used in the pediatric clinic because it is predictive of the outcome and adaptive skills of children with neurodevelopmental symptoms^15^.

Due to statistical power, most studies have repeatedly analyzed a small set of the most frequently recurrent CNVs (population frequency >1/10,000)^16–18^, which collectively affect only approximately 2% of the coding genome^19^. As a result, our understanding of gene functions sensitive to gene dosage is highly biased. However, the vast majority of CNVs affecting neurodevelopmental and cognitive ability are ultra-rare (<1/10,000)^17^. and have been implicated based on their size, and gene content using burden analyses^12,19–22^. Such CNVs cover a large proportion of the coding genome and remain difficult to study individually with currently available sample sizes. Beyond CNVs, more generally, our understanding of gene-disrupting variants associated with cognitive ability and neurodevelopmental disorders stems from approximately 200 genes disrupted by *de novo* variants^4,23^. Their functions are enriched in chromatin and transcription regulation, regulation of nervous system development, central nervous system neuron differentiation, and regulation of synapse structure and activity^4,23^. It is unclear if these functions are most representative of cognitive ability or genetic fitness.

### Knowledge gap

Overall, it has been difficult to investigate the broad landscape of ultra-rare CNVs potentially involved in neurodevelopmental traits, such as cognitive ability. As a result, we have a limited understanding of the full range of gene-dosage-sensitive biological processes linked to cognitive ability.

To circumvent the issue of power, research groups, including ours, have implemented alternative approaches aggregating rare variants disrupting genes with similar constraint scores in order to perform “constraint burden” association studies^11,12,19–21,24^. These burden analyses showed that genes with increasing intolerance to haploinsufficiency were associated with increasing effect sizes on cognitive ability and risk for psychiatric illnesses, such as ASD, schizophrenia, and bipolar disorder^19,21^. Similarly, studies have developed methods to aggregate common variants^25^, demonstrating that a robust association with a condition (e.g., ASD) can be established at the group level when individual single nucleotide polymorphisms (SNPs) do not meet genome-wide criteria for association.

In this study, we aimed to investigate the full range of gene-dosage-sensitive biological processes linked to cognitive ability. To this mean, we analyzed all CNVs >50 kilobases in 258k individuals across 6 cohorts from the general population. The CNV-level GWAS identified the first CNV associated with higher cognitive ability. We then performed functional-burden associations with cognitive ability across 6,502 gene-sets corresponding to gene functions at the tissue, cell type, and molecular levels. Findings reveal that most biological functions have preferential effects on cognitive ability when either deleted or duplicated. As a result, we observed a negative correlation between the effects of deletions and duplications across all levels of biological observation independently of intolerance to haploinsufficiency.

## Methods and materials

### Cohorts

We analyzed 258,292 individuals from six general population cohorts^26–31^, which can be further divided into 9 sub-cohorts based on cognitive assessment (Table1). Three additional autism cohorts^32–34^ were only used for sensitivity analyses (Supplementary table 1, figure 1, and Methods). Each cohort received approval from their local institutional review boards. Parents/guardians and adult participants gave written informed consent, and minors gave assent.

**Table 1.** Cohort descriptions. Analyses were performed (after QC) in 258,292 individuals from 6 general population cohorts. SYS: Saguenay Youth Study, CaG: CARTaGENE, LBC1936: Lothian Birth Cohort 1936; N=number of individuals remaining for analysis after quality control. The mean and Standard Deviation (SD) for g-factor slightly deviate from 0 and 1 in some cohorts since they were computed on all available data (before the exclusion of some individuals for poor quality array) and summarized here only for individuals included in the analyses. cognitive measures used to compute g-factors are detailed in Supplementary Table 1. Autism cohorts used only for sensitivity analyses are detailed in Supplementary Table 1.

| Cohort |  | N= | Ancestry |  | Gender |  | Age (year) |  | Cognitive ability assessments |
| --- | --- | --- | --- | --- | --- | --- | --- | --- | --- |
|  |  |  | EUR | Others | F | M | Mean | SD |  |
| Unselected<br>(n=258,292) | CaG | 2,589 | 2,472 | 117 | 1,375 | 1,214 | 53.943 | 7.845 | g-factor |
|  | G-Scot | 13,715 | 13,672 | 43 | 8,081 | 5,634 | 46.730 | 14.996 | g-factor |
|  | Imagen | 1,744 | 1,624 | 120 | 891 | 853 | 14.450 | 0.366 | WISC-IV |
|  | LBC1936 | 503 | 500 | 3 | 246 | 257 | 69.825 | 0.829 | Moray House Test<sup>65</sup> |
|  | SYS | 1,565 | 1,561 | 4 | 824 | 742 | 28.177 | 17.098 | WISC-III (g-factor) |
|  | UKBB | 73,882 | 71,364 | 2,518 | 39,317 | 34,565 | 60.022 | 8.959 | g-factor<sup>66</sup> |
|  |  | 62,080 | 60,484 | 1,596 | 34,335 | 27,745 | 62.083 | 7.663 | g-factor (online) |
|  |  | 88,441 | 80,427 | 8,014 | 47,789 | 40,652 | 58.139 | 8.304 | FI |
|  |  | 13,773 | 13,458 | 315 | 8,284 | 5,489 | 64.185 | 7.685 | FI (online) |

**Figure 1.**
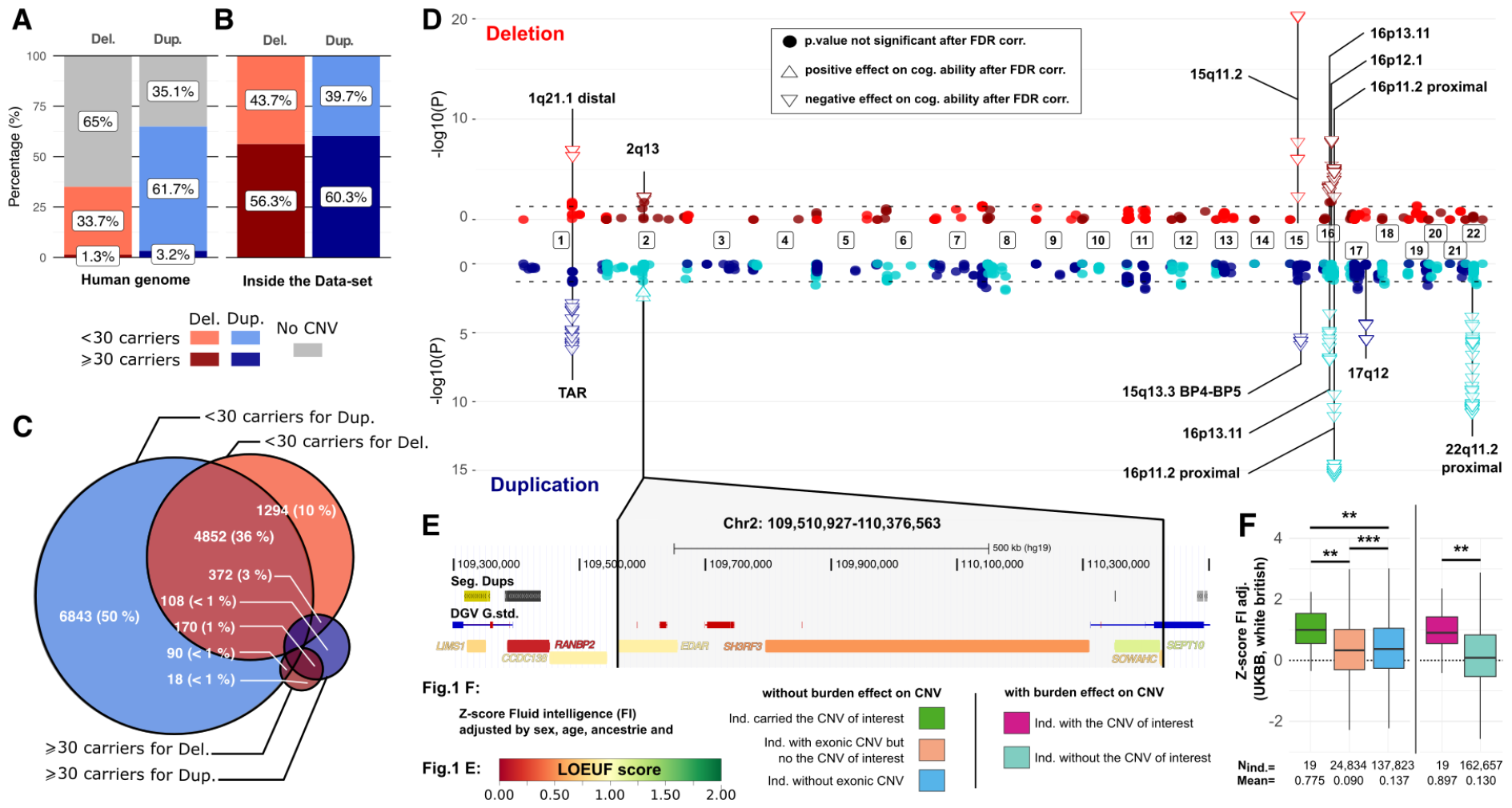
CNV GWAS on general cognitive abilities at the gene level. (A) Proportion of genes deleted (red) or duplicated (blue) at least once in the dataset, among all genes in the human genome. Deleted or duplicated genes observed in less than 30 carriers (light color) and 30 or more (dark color) as well the proportion of genes not observed in any CNV (grey),(n=13,747). (B) The majority of deleted or duplicated genes were observed in more than 30 carriers. (C) Venn diagram illustrating the overlap between genes deleted and-or duplicated at least once in the dataset. (D) The Miami plot illustrates the −log10 transformed p-value of the association with cognitive ability for each gene included in deletions (red) at the top, and duplications (blue) at the bottom, along the genome. Adjacent chromosomes are shown in alternating light and dark colors. Triangles represent significant genes after FDR correction, while circles represent non-significant genes. The direction of the triangle indicates the effect size. The dash line represents the nominal significant pvalue threshold. (E) A specific duplication, chr2:109,510,927-110,376,563 (including *EDAR*, LOEUF=0.91; *SH3RF3*, LOEUF=0.53; *SEPT10*, LOEUF = 1.17; *SOWAHC*, LOEUF=0.77), exhibited a previously unobserved positive effect on cognitive ability in the CNV GWAS. (F) To further investigate this positive effect, we conducted a post hoc analysis using a two-sided t-test on a homogeneous cohort with consistent technology, ancestry, and phenotype, aiming to eliminate biases. Our focus was specifically on individuals of white British ethnicity in the UK Biobank (UKBB) with adjusted fluid intelligence. In the left part of the analysis (F), individuals were categorized into three groups: carriers of the CNV of interest (green), non-carriers of this specific CNV but carrying other exonic CNVs (light orange), and non-carriers of any exonic CNV (blue). The t-tests were performed on fluid intelligence (FI) adjusted for sex, 1 to 10 PC for ancestry, and age. In the right part of the analysis (F), two groups were defined: carriers of the CNV of interest (dark pink) and non-carriers (light blue). The t-tests were conducted on fluid intelligence adjusted for sex, ancestry, age, and the burden of 1/LOEUF for deletions and duplications. For the duplication of chr2:109,510,927-110,376,563 observed in the CNV GWAS (F), the carriers exhibited significantly higher cognitive ability measures compared to both other groups. Furthermore, when we weighted the cognitive ability by the burden of 1/LOEUF for deletions and duplications, a positive effect was also observed among carriers of the CNV of interest.

### Measures of cognitive ability

General cognitive ability was measured by either non-verbal intelligence quotient (NVIQ), fluid intelligence (fluid intelligence questions), or general intelligence factor (g-factor)^15^. Measures of cognitive ability were z-scored within each cohort based on sex and age (table 1, Supplementary table 1 and 2; Supplementary Methods).

### Array processing and genotyping

All arrays used for genotyping and CNV calls are detailed (Supplementary table 1 and Supplementary method)^12^. We used pLink^35^ to check for duplicates, sex, and family relationships for each participant using a standard pipeline^11,12^. Ancestry principal components and categories were computed using KING (www.kingrelatedness.com) with the 1000 Genomes Project^11,12^ as the reference population.

### CNV calling and quality control

For CNV calling, we used PennCNV^36^ and QuantiSNP^37^. For array and CNV quality control, we applied standard filtering parameters (Supplementary Methods, https://martineaujeanlouis.github.io/MIND-GENESPARALLELCNV/)^11,12^. For the MSSNG dataset^34^, we used published CNVs called on whole genome sequencing^38^. To harmonize CNV calls across arrays with different probe densities, we only considered CNVs encompassing at least 10 probes present on all array technologies. We excluded from the analyses all individuals carrying a CNV ≥10Mb or a mosaic CNV.

### Gene annotation

We annotated autosomal CNVs using bedtools (*https://bedtools.readthedocs.io/en/latest/*) with Gencode V19 (hg19, Ensembl gene name, *https://grch37.ensembl.org/index.html*).

### LOEUF-based and function-based gene sets

We defined 38 overlapping gene-sets based on LOEUF values (loss-of-function observed/expected upper bound fraction, gnomAD version 2.1.1)^39^ (Supplementary methods). LOEUF range from 0 to 2, and values below 0.35 are suggestive of intolerance.

We defined 269 gene-sets based on relative gene expression (Z-score >1SD) in 13 adult^40,41^ and 16 fetal^42^ brain cell types^43^, as well as bulk tissue from 215 brain regions (Human Protein Atlas, HPA v.22)^44^ and 25 non-brain organs (GTEx^44,45^, Supplementary table 3). As a sensitivity analysis, we defined the same gene-sets based on a previously published “Top Decile Expression Proportion” (TDEP)^46^ method. The former and the latter methods favour relative and specific expression respectively. Both methods exclude 1,370 and 5,369 genes that are not assigned to any tissue in GTEx. We also used 6,233 functional gene-sets based on 6,130 GOterms^47,48^ (Ensembl v.109, April 2023), and 103 Synapse ontology terms (SynGO^49^). Throughout this study, we only considered gene-sets meeting the following 3 criteria: i) those with more than 10 genes, ii) those disrupted by ≥ 30 CNV carriers, and iii) those with at least 20% of their genes affected by CNVs.

### Statistical analyses

Analyses were performed using R version 4.0.1(http://www.R-project.org.), with “meta”(*https://cran.r-project.org/web/packages/meta/index.html*) and “metafor”(*https://cran.r-project.org/web/packages/metafor/index.html*) packages for meta-analyses. Python 3.10.2(https://www.python.org) with “scipy 1.11.2”(*https://pypi.org/project/scipy/*), “statsmodels 0.13.5” (https://www.statsmodels.org) and “word-cloud 1.9.2” (https://amueller.github.io/word_cloud).

### LOEUF-based and Function-based burden association tests

To estimate the effect on cognitive ability of gene-sets (and their corresponding biological functions or LOEUF categories), we adapted a previously published model^12^. We performed a linear model for each of the 38 LOEUF gene categories and each of the 6502 functional gene-sets. The outcome was cognitive ability measured in each individual. The explanatory variable was the sum of genes fully encompassed in a CNV for a gene-set of interest (Figure 2A). Since CNVs are multigenic, the effect size estimated for a given gene set may be inflated. Therefore, all models were adjusted for the total number of genes within the CNV but not members of the gene set of interest. These latter genes were categorized into three variables: ID genes (only for Function-based gene-sets), genes with LOEUF<1, and genes with LOEUF≥1. Other covariates included ancestry (10 PCs), age, and sex. Models were computed for deletions or duplications, separately (Supplementary method).

**Figure 2.**
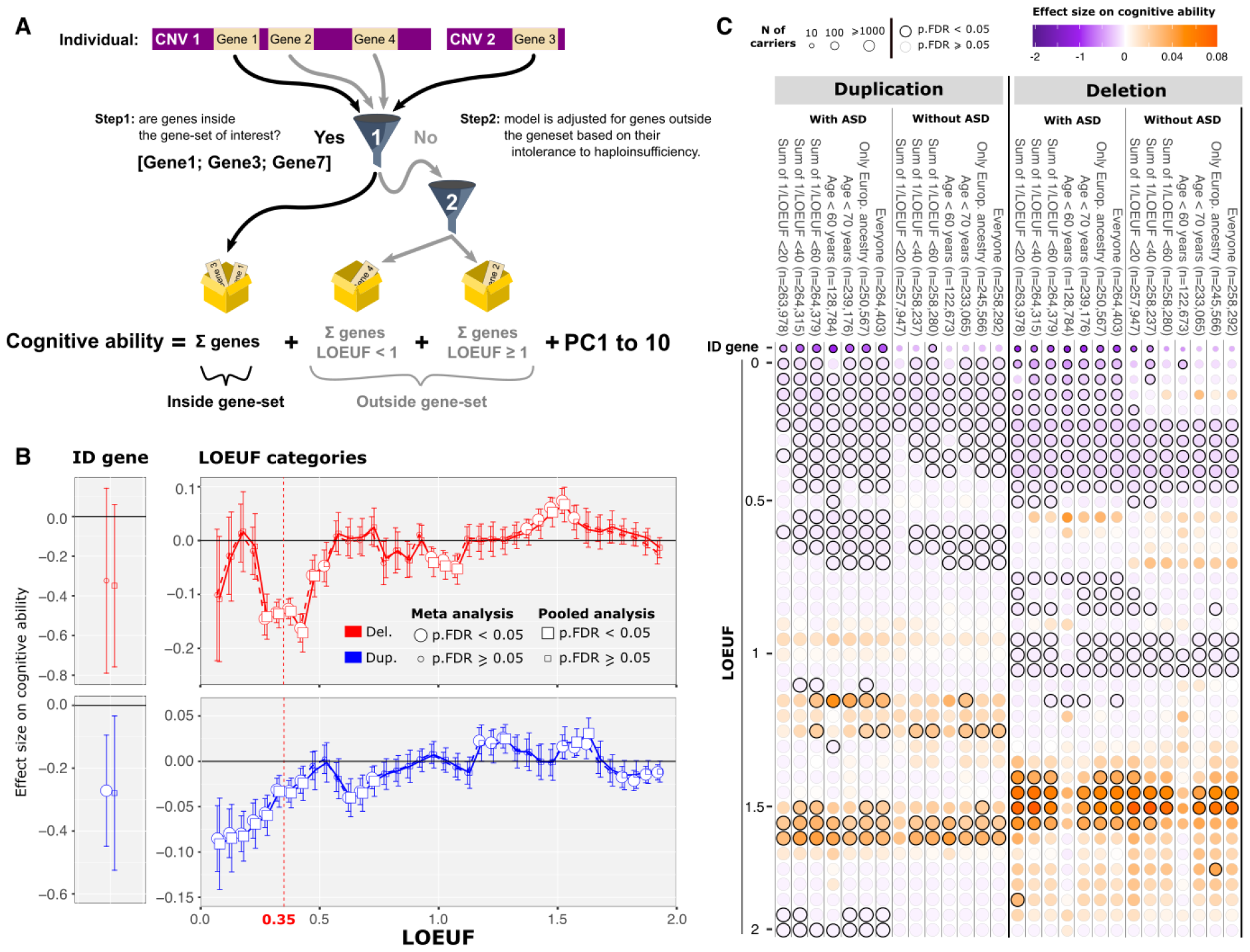
Effect sizes of autosomal coding genes on general cognitive abilities based on their LOEUF values. (A) The functional burden test is a linear model estimating the mean effect size of all CNVs fully encompassing genes assigned to a biological function of interest. Because many CNVs are multigenic, the model is adjusted for genes included in a CNV but not assigned to the biological function of interest. (B) Sliding window (Supplementary method statistical model 2) estimating the effect size on cognitive ability of deletions (top) and duplications (bottom) for 38 LOEUF categories (we slide a window of size 0.15 LOEUF units, in increments of 0.05 units thereby creating 38 categories across the range of LOEUF values) and definitive ID genes curated by ClinGen. Estimates were computed using a meta-analysis (circles) as well as a pooled dataset (squares). The red dashed line defines intolerant genes (LOEUF<0.35). (C) Heatmap showing the effect size (color scale) on cognitive ability of deletions and duplications across a range of sensitivity analyses removing non-europeans, older participants, large multigenic CNVs (those with a sum of 1/LOEUF >60, >40, >20 corresponding to values of well known recurrent CNVs: 22q11.2, 16p11.2 and TAR respectively) as well as adding a neurodevelopmental dataset (autism spectrum disorder). All estimates were computed on the pooled dataset.

### tagDS

Previous publications have reported that effect size on cognitive ability is U-shaped. Effects of deletions are 2–3 times higher than duplications^12^. To test whether the deletion/duplication effect size ratio deviates from the expected average ratio (specifically 2.4 in our dataset), we developed the trait-associated gene-dosage sensitivity score (tagDS) (Supplementary method). tagDS was normalized (Z-scored) based on a null distribution of tagDS values. The latter distribution was obtained using effect sizes on cognitive ability computed for deletions and duplications across 570,000 randomly sampled gene-sets. A tagDS of 0 suggests that the deletion/duplication effect-size ratio is equal to the expected ratio. A gene-set with a tagDS > 2 indicates that its deletion/duplication effect-size ratio on cognitive ability is beyond 2 standard deviations of the null distribution (i.e., larger effect sizes are biased towards deletions).

## Results

### Gene dosage may be associated with higher cognitive ability

We observed that 15.6% of the 258,292 individuals included in the analysis carried at least one autosomal CNV fully encompassing one or more coding genes. Among 18,451 autosomal coding genes with LOEUF values, 35% and 64.9% were fully encompassed in one or more deletion and duplication, respectively (75% across CNVs), with 40% observed in both deletions and duplications (Figure 1A, B, C). Most genes were in ultra-rare CNVs (<1/10,000) with fewer than 30 carriers (Figure 1C). We computed a linear regression model (gene-level GWAS, Supplementary method statistical model 1) on general cognitive ability for 241 and 596 genes covered by at least 30 deletions or duplications, respectively, in 258,292 individuals from the general population (Figure 1D). We identified 9 deletions encompassing a total of 69 genes and 10 duplications encompassing a total of 123 genes with previously published negative effects (Supplementary table 4) that persisted when we conducted a meta-analysis across 9 sub-cohorts defined by cognitive assessments (Table 1, Supplementary figure 2). We identified a novel association between a duplication at 2q12.3, and positive effects on cognitive ability (Figure 1E). This duplication observed in 36 individuals included 4 genes with a LOEUF≥0.35 (*EDAR, SH3RF3, SEPT10, SOWAHC*) and was observed at a similar frequency (1 to 2 in 10,000) across cohorts (Fisher’s exact p-value_FDR_ >0.05). Results were not related to ancestry, array platform, or cognitive assessment methods (Figure 1F). The positive effect remained significant when comparing 2q12.3 duplication carriers to individuals without any CNVs.

### A large proportion of intolerant and tolerant genes modulate cognitive ability

Even with the current large sample sizes, CNVs cover only 3-4% of coding genes in our previous analyses. We know that a much larger proportion of the coding genome is involved in cognitive ability^12^, we used burden association methods to explore all coding CNVs. We partitioned genes into 38 sets based on overlapping LOEUF categories and an ID gene-set(as defined by ClinGen Supplementary table 5), then calculated their 39 mean burden effect sizes using linear models adjusted for CNVs outside each category to prevent effect inflation from multigenic CNVs(c.f. methods, Supplementary method statistical model 2, Figure 2A). The 39 estimates provided by the meta-analysis across the 9 sub-cohorts were not different from those provided by aggregating these datasets (Figure 2B, Supplementary table 6 and 7). Therefore, all subsequent analyses were performed on the aggregated dataset. The effects of deletions were, on average, 2.4-fold higher than duplications, and we observed a positive correlation between the effect sizes of deletions and duplications across LOEUF categories (Spearman’s r=0.5, p_permutation_=0.02; Supplementary figure 3). Negative effects on cognitive abilities were observed in 8 and 11 non-tolerant categories (LOEUF<1) for deletions and duplication, respectively. More intolerant the LOEUF category was, the more negative the effect size, with the ID gene-set having the largest effects. Of note, 2 and 3 categories showed positive effects for deletions and duplications, respectively. In other words, the effect sizes of these categories were significantly higher than the average effect of gene categories used to adjust the model. Sensitivity analyses showed no biases related to ancestry, large multigenic CNVs, or low-quality control scores (Figure 2C, Supplementary figure 4). Effect sizes of intolerant genes were higher when removing older age groups (>60 years old) (Figure 2C). Because the most intolerant CNVs are depleted in the general population, we included 3 ASD cohorts in a sensitivity analysis. This resulted in larger effects and smaller p-values for highly intolerant LOEUF categories without changing the effects of other categories ≥0.35 (Figure 2C).

### Negative correlation between deletion and duplication effects on cognitive ability across brain regions

Previously published functional enrichment analyses^50,51^ have focused on recurrent CNVs. We, therefore, developed a functional burden test to systematically investigate gene functions that may underlie the pervasive association between CNVs (too rare to reach individual association) and cognitive ability. We first focused on gene sets assigned to 215 adult brain regions. 91 and 94 were associated with cognitive ability when deleted and duplicated, respectively, and only 25 were impacted by both(c.f. methods, Figure 3A). This suggests that cognitive ability is preferentially affected by one CNV type, depending on the brain region. These preferential effects were supported by the negative correlation observed between the effect sizes of CNV type across all brain regions (Spearman’s r=-0.43, p_permutation_=9×10^−3^; Figure 3B). Stratifying these brain gene-sets into 3 independent LOEUF categories provided the same negative correlations (Figure 3C). Sensitivity analysis showed that the negative correlation was not due to unbalanced power between deletions and duplications or the relative expression threshold used to define gene-sets (Supplementary figure 5). To further investigate gene-sets driving this negative correlation, we developed the “trait-associated gene-dosage sensitivity score” (tagDS), a normalized value reflecting whether a gene-set shows preferential effects on cognitive ability when either deleted or duplicated (cf method, Figure 3D). TagDS indicated that cerebral cortex gene-sets affected cognitive ability preferentially when duplicated, while the opposite was observed for non-cortical (subcortical and midbrain) gene-sets and deletions (Figure 3A, Mann-Whitney p_permutation_=1×10^−15^). The same cortical/non-cortical gene dosage sensitivity was also observed when removing genes with low tissue specificity (Supplementary figure 5)^44^.

**Figure 3:**
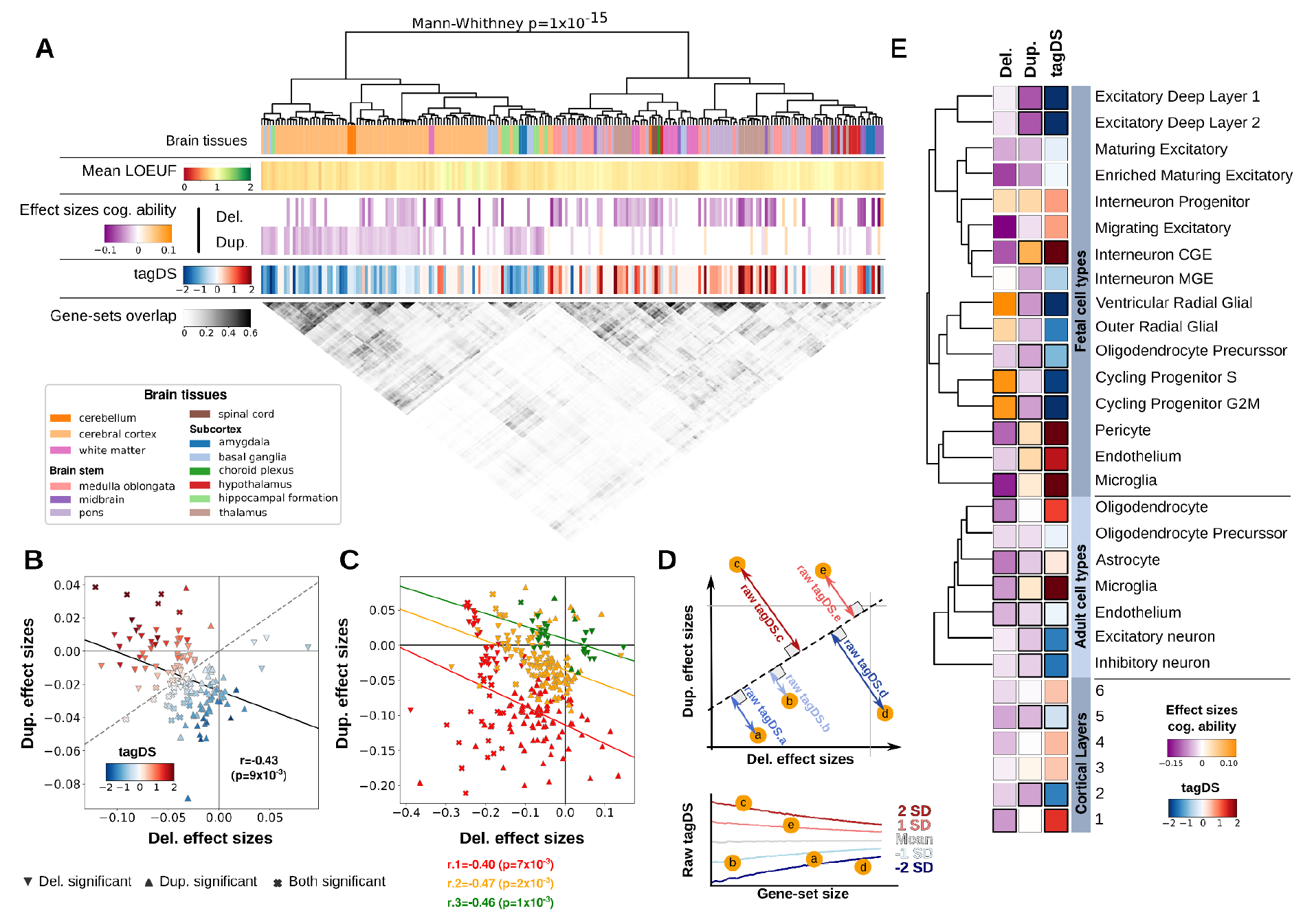
Effects on cognitive ability of genes assigned to brain regions and cell types. (A) Effects sizes on cognitive ability of gene sets assigned to 215 brain tissues/regions. Brain regions are color-coded and clustered (first row, Wards’method^67^) based on the level of overlap (gray matrix) between their corresponding gene-sets. The average LOEUF value for each gene-sets is color-coded in the 2nd row. The mean effect sizes, on cognitive ability, of genes assigned to each brain region, are coded for deletions (3rd row) and duplications (4th row). TagDS values are represented in the 5th row. (B) Spearman correlation (black line) between the effect sizes of deletions and duplications across all gene-sets with FDR significant effects on cognitive ability for either deletions (downward triangle), duplications (upward triangle), or both (cross). P-values were obtained from permutations to account for the partial overlap between gene-sets. Gene-sets are color-coded based on their tagDS. The dashed line represents the average exome-wide duplication/deletion effect-sizes ratio. (C) The same negative correlations between deletion and duplication were observed across 3 independent LOEUF groups: <0.35 (intolerant; red), [0.35, 1.0[ (moderately intolerant; orange), and [1.0, 2.0] (tolerant; green). (D) Raw tagDS is the Euclidean distance to the whole-genome ratio of effect sizes. TagDS is normalized following the null distribution of random gene-sets of identical size. (E) Effect size of deletions and duplications encompassing genes assigned to 6 cortical layers, 7 adult brain cell types, and 16 fetal brain cell types. Clustering was calculated on the level of overlap between cell-type gene-sets (Wards’method^67^). Purple and orange represent negative and positive effects on cognitive ability, respectively. Black edges indicate significant effects.

At the microstructure and cell-type level (6 cortical layers, 7 adult, and 16 fetal brain cell types based on their normalized gene expression, c.f. methods), we observed the same negative correlation (r=-0.70, p_permutation_<1×10^−3^; Supplementary figure 6). The largest effects for deletions and duplications were observed in gene-sets assigned to fetal cell types. Deletions and duplications, respectively, showed preferential effects in non-neuronal (endothelial, glia) and neuronal (excitatory) cell types (Figure 3E).

### Genes involved in non-brain tissues affect cognitive ability

There is a growing interest in whole-body health comorbidities among individuals with neurodevelopmental and psychiatric conditions as well as CNVs affecting cognition^18,52^. We, therefore, asked if CNVs affecting genes assigned to non-brain tissues were also associated with cognitive ability. We used 37 gene-sets, defined by relative expression in 12 brain and 25 non-brain tissues (>1SD, Figure 4A). Many non-brain gene-sets showed effect sizes (Figure 4B) of similar magnitude to those observed for regional brain gene-sets. This was not explained by the level of overlap between brain and non-brain gene-sets (Figure 4A, B). We observe the same pattern of deletion-duplication negative correlation independently of the gene-set definitions (r=-0.64 p_permutation_<1×10^−3^; Figure 4C; Supplementary figure 7). To understand how gene-set definitions influence these results, we first removed 8,194 genes with low-tissue specificity assigned to multiple gene-sets. The resulting effect sizes were correlated with the initial estimates (r=0.57, Supplementary figure 8). In fact, genes assigned to multiple tissues show higher intolerance (LOEUF) compared to tissue-specific genes (p=1×10^−11^ to 3×10^−161^, figure 4D, Supplementary figure 9). To further investigate the impact on results of gene-sets definition, we tested 37 previously published gene-sets assigned to 37 GTEx tissues computed by the TDEP method (proportional gene expression)^46^. This method, which emphasizes specificity, excludes 5,454 genes of which 1,586 and 696 are respectively moderately (LOEUF [0.35, 1[) and highly intolerant (LOEUF<0.35; Supplementary figure 9). Effect sizes were well correlated with our analysis excluding LTS genes (r=0.76), but TDEP gene-sets were unable to detect any effect for deletions across all tissues (Supplementary figure 8).

**Figure 4:**
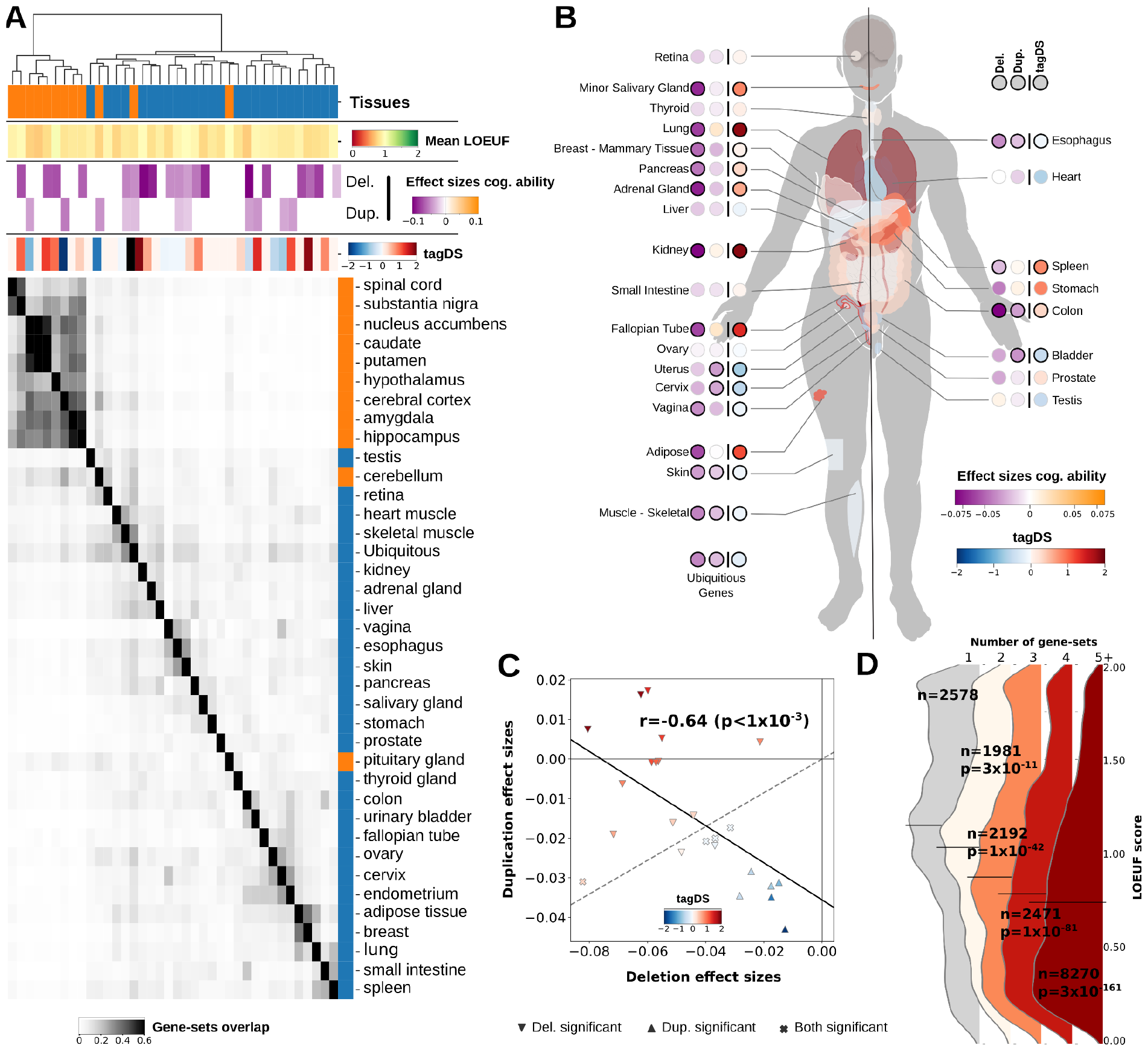
Effects on cognitive ability of CNVs affecting genes implicated in brain and non-brain tissues. (A) We defined 37 gene-sets based on z-scored expression > 1SD. Expression of each gene was normalized across 37 tissues provided by GTEx. Gene sets were clustered (orange for brain tissues and blue for non-brain tissues) based on their overlap which is shown in the grayscale matrix. High overlap was observed between brain gene-sets (Wards’method^67^), and much lower overlap was present across non-brain tissues and between brain and non-brain tissue. The mean LOEUF of each gene-set is color-coded in row 2. Effect sizes on cognitive ability and tagDS across tissues are color-coded in the 3rd row as well as in the body map (B), adapted from GTEx. Genes with low tissue specificity were defined by the Human Protein Atlas. (C) Spearman correlation (black line) between the effect sizes of deletions and duplications on cognitive ability. Downward and upward triangles, as well as crosses, represent significant effects for deletions, duplications and both respectively. Gene-sets are color-coded based on their tagDS. (D) Distribution of LOEUF values for genes assigned to 1, 2, 3, 4, or ≥5 gene-sets. The 5 groups are non-overlapping. Black lines represent the median LOEUF score for each group. The p-values are provided by the Mann-Whitney test between each category and the genes assigned to one gene-set.

### The effects of deletion and duplication on cognitive ability are negatively correlated across all levels of biological observations

We asked if the deletion-duplication negative correlations observed for tissue-level gene-sets were also present at the molecular and cellular components levels. We first investigated 293 synaptic gene ontologies using SynGO^49^. We observed that post-synaptic genes showed the largest negative effects on cognitive ability when deleted, and in contrast, pre-synaptic genes when duplicated (Figure 5A, Supplementary figure 10). As a result, the effects of the 2 opposing CNVs were negatively correlation across SynGO terms (r=-0.39, p_permutation_=1×10^−3^; Supplementary figure 11).

**Figure 5.**
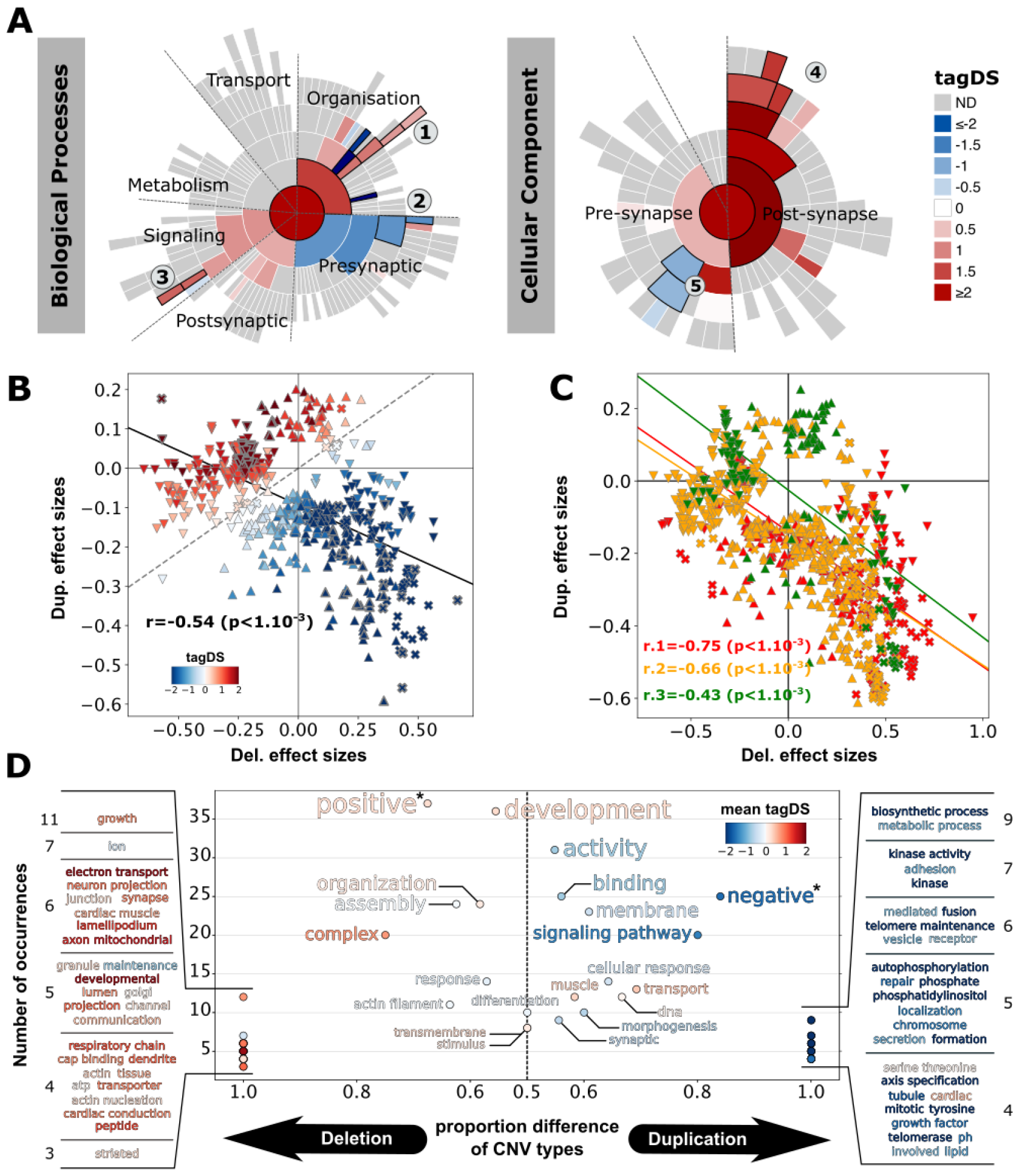
Effects on cognitive ability of gene sets based on gene ontologies. (A) Effect sizes of synaptic molecular functions and cellular component gene-sets as defined by SynGO^49^ on cognitive ability (More details in the supplementary figure 10). Blue and red represent negative and positive cognitive ability tagDS, respectively. Ontologies with black edges indicate significant effects (FDR). The results are shown only for SynGO terms with more than 10 genes, observed at least 30 times in our dataset, and with a coverage greater than 20%. Note panel (A): 1) Regulation of modification of postsynaptic actin cytoskeleton, 2) Regulation of calcium-dependent activation of synaptic vesicle fusion, 3) Presynaptic modulation of chemical synaptic transmission, 4) Integral component of postsynaptic density membrane, 5) Synaptic vesicle membrane. (B) There is a negative correlation (Spearman) between the effect sizes of deletions and duplication across 601 GO-terms. (C) The same deletion-duplication negative correlation was observed across 3 independent LOEUF groups (intolerant <0.35: red, moderately intolerant [0.35, 1.0[: orange, tolerant [1.0, 2.0]: green). (D) We adapted the word cloud package, which groups GOTerms based on shared terminology. Y-axis: Sum of associations of each word with significant deleted and duplicated GOTerms. X-axis: Proportion of significant GOTerms for a given CNV type used for the association. “Positive” and “Negative” refer to “Positive regulation” and “Negative regulation”.

We extended our analysis to 6,130 GO-terms (and corresponding gene-sets), 5.0% and 3.5% of the GO-terms had an effect size on cognitive ability for deletions and duplications, respectively. A minority (0.7%) of GO-terms showed significant effects for both. We observed again a deletion-duplication negative correlation across GO-terms effect sizes (r=-0.54, p_permutation_<1×10^−3^; Figure 5B), which remained significant across three independent levels of LOEUF stratification (Figure 5C). We asked if tagDS was similar to pHI (probability of haploinsufficiency) and pTS (probability of triplosensitivity), 2 previously published metrics that are highly correlated with each other (0.78) and with LOEUF (r=0.90 and 0.77, respectively). tagDS was unrelated to pHI and pTS scores^50^ across GO-terms (Supplementary figure 15). Finally, several GO-terms, such as neuronal, synaptic, and cardiac functions, showed preferential effects when deleted, while the opposite was observed for cellular response functions, transport, metabolic processes, and signaling pathways. Furthermore, “positive regulation” GO-terms were more sensitive to deletions, while “negative regulation” terms showed preferential effects when duplicated (Figure 5D, Supplementary figure 12).

## Discussion

In this large-scale CNV-GWAS on cognitive ability, we identified a duplication at 2q12.3 that is associated with higher cognitive ability. Although our sample size limited the discovery of new genome-wide signals at the variant-level, we developed a functional burden association test that allowed us to simultaneously test the contribution of all ultra rare CNVs (covering 75% of the coding genome) and their function to cognitive ability. Constraint (LOEUF), and functional burden analyses revealed that a substantial portion of the coding genome was associated with cognitive ability when deleted or duplicated. We also demonstrated that genes involved in a broad array of biological functions show preferential effects on cognitive ability when either deleted or duplicated. The latter was quantified by negative correlations between deletion and duplication effect sizes and by tagDS, a new normalized metric that assesses sensitivity to either deletions or duplications. We also show that genes assigned to non-brain tissues affected this “brain-centric” trait.

We identify, to our knowledge, the first CNV associated with higher cognitive ability. The 865kb duplication (population frequency ∼1/7200), which includes *EDAR, SH3RF3, SEPT10, SOWAHC*, had not been previously associated with any trait or condition. Publications have identified associations between SNPs within this locus and 58 traits, including brain morphology^54–56^, schizophrenia^57,58^, Alzheimer’s disease^59^, and neuroinflammatory biomarkers^60^ (Supplementary table 8). An excess of *SEPT10 de novo* missense mutations have been reported in neurodevelopmental disorders^4^. Given that the median age of our dataset is 60.7 years, it is possible that this duplication may be associated with a neuroprotective effect. We suspect that many more CNVs associated with higher cognitive ability will be identified in the future as sample sizes increase. Our functional burden method identified gene-sets with positive effects on cognitive ability. Determining whether these gene-sets truly increase cognitive ability or, instead, show smaller effects than the mean effect used to adjust for multigenic CNVs will require larger samples with data on CNVs disrupting single genes. Overall, results suggest that gene dosage may be associated with higher IQ but most effects are masked by the multigenic nature of CNVs.

It has been challenging to evaluate **haploinsufficiency** and **triplosensitivity**. We show that tagDS for cognitive ability is orthogonal to genetic constraint, as well as previously published pHI and pTS measures. The tagDS highlights sensitivity to either deletions or duplications across gene functions from macroscopic (cortical versus non-cortical tissue) to microscopic (pre versus postsynaptic genes and positive versus negative regulation) levels of observation.

**Genetic covariance** has almost exclusively been computed using common variants to investigate the genetic overlap between traits. While genetic covariance using rare variants are understudied due to a lack of statistical power, a recent study^61^ aggregating rare variants at the gene level showed that the genetic correlation between protein loss-of-function and damaging missense variants associated with the same trait was on average 0.64 (with some correlations < 0.5) implying that different classes of variants in the same genes may show different phenotypic effects.

In our study, we show that two classes of variants with opposing molecular consequences have negatively correlated phenotypic effects. This negative correlation was observed regardless of whether CNVs were aggregated based on their function in tissues, cell types, or GOterms. This suggests that associating genes to traits or diseases is highly dependent on the class of genetic variants. Whether this negative correlation generalizes to other phenotypic traits is unknown.

There has been growing interest in the relationship between mental health and whole-body multi-morbidities. This is exemplified by the correlation between cognitive ability, medical conditions, such as coronary artery disease^15,62^ and longevity^62,63^. Recent studies also showed that poor physical health was more pronounced in neuropsychiatric illness than poor brain health^52^. In the current study, genes preferentially expressed in many non-brain organs show effects on cognition similar to those observed for brain tissue. The latter could not be explained by the level of overlap between brain and non-brain gene-sets. However, our results suggest a trade-off between genesets including intolerant genes those emphasizing tissue specificity. In other words, genes with lower tissue specificity and higher pleiotropy tend to have lower LOEUF values and therefore larger effect sizes on cognitive ability. Other interpretations include: i) gene-disrupting variants can alter non-brain organs, which in turn alter brain function due to suboptimal support; and ii) cognition is an embodied multi-organ trait includes both brain and non-brain organs. A whole-body contribution exists for other cognitive-modulating traits such as sleep (thought to be for and by the brain), which is also regulated by peripheral tissue^64^.

The main limitation of this study is the use of gene-sets, which were either defined on the basis of well-established ontologies or using a “relative method” based on normalized expression values. In the latter approach, we chose thresholds that may have influenced our results. Multiple sensitivity analyses demonstrated that changing the threshold (and, therefore, the size of the gene-set) did not influence our main findings. Expression profiles vary across space, cell-types, and time for a given tissue. Our gene-sets could not explore all of these aspects. Larger studies will be required to increase the granularity of these functional burdens on association tests.

In conclusion, our study demonstrated, for the first time, positive effects of a CNV on cognitive abilities. Our analysis presents a new statistical approach for partitioned rare variant association and uncovers many gene functions that are preferentially sensitive to either deletions or duplications. Computing tagDS for other complex traits will help understand whether sensitivity to gene dosage is trait-dependent.

## Funding/Support

This research was enabled by support provided by Calcul Quebec (http://www.calculquebec.ca) and Compute Canada (http://www.computecanada.ca). Sebastien Jacquemont is a recipient of a Canada Research Chair in neurodevelopmental disorders, and a chair from the Jeanne et Jean Louis Levesque Foundation. This work is supported by a grant from the Brain Canada Multi-Investigator initiative and CIHR grant 159734 (Sebastien Jacquemont, Tomas Paus). The Canadian Institutes of Health Research and the Heart and Stroke Foundation of Canada fund the Saguenay Youth Study (SYS). SYS was funded by the Canadian Institutes of Health Research (Tomas Paus, Zdenka Pausova) and the Heart and Stroke Foundation of Canada (Zdenka Pausova). Funding for the project was provided by the Wellcome Trust. This work was also supported by an NIH award U01 MH119690 granted to Laura Almasy, Sebastien Jacquemont and David Glahn and U01 MH119739. The authors wish to acknowledge the resources of MSSNG (www.mss.ng), Autism Speaks and The Centre for Applied Genomics at The Hospital for Sick Children, Toronto, Canada. We also thank the participating families for their time and contributions to this database, as well as the generosity of the donors who supported this program. We thank the coordinators and staff at the SCC sites. We are grateful to all of the families at the participating SSC sites and the principal investigators (A. Beaudet, M.D., R. Bernier, Ph.D., J. Constantino, M.D., E. Cook, M.D., E. Fombonne, M.D., D. Geschwind, M.D., Ph.D., R. Goin-Kochel, Ph.D., E. Hanson, Ph.D., D. Grice, M.D., A. Klin, Ph.D., D. Ledbetter, Ph.D., C. Lord, Ph.D., C. Martin, Ph.D., D. Martin, M.D., Ph.D., R. Maxim, M.D., J. Miles, M.D., Ph.D., O. Ousley, Ph.D., K. Pelphrey, Ph.D., B. Peterson, M.D., J. Piggot, M.D., C. Saulnier, Ph.D., M. State, M.D., Ph.D., W. Stone, Ph.D., J. Sutcliffe, Ph.D., C. Walsh, M.D., Ph.D., Z. Warren, Ph.D., and E. Wijsman, Ph.D.). We appreciate obtaining access to phenotypic data on SFARI base. LBC1936 is supported by the Biotechnology and Biological Sciences Research Council, and the Economic and Social Research Council [BB/W008793/1] (which supports SEH), Age UK (Disconnected Mind project), the Milton Damerel Trust, and the University of Edinburgh. SRC is supported by a Sir Henry Dale Fellowship jointly funded by Wellcome and the Royal Society [221890/Z/20/Z]. Genotyping was funded by the BBSRC (BB/F019394/1).

## Role of the Funder/Sponsor

The funder had no role in the design and conduct of the study; collection, management, analysis, or interpretation of the data; preparation, review, or approval of the manuscript; or decision to submit the manuscript for publication.

## Footnotes

Conflict of interest: The authors declare that they have no conflicts of interest.

## Notes

### Competing Interest Statement

The authors have declared no competing interest.

