## Supplementary method and Figures for "Effects of gene dosage on cognitive ability: A function-based association study across brain and non-brain processes"

### Supplementary methods

*General populations*

In this study, we included five cohorts from the general population previously pooled and studied in Huguet et al 2021[1]. In addition to these cohorts previously analyzed and studied, we added 238,176 individuals from the UK Biobank (UKBB) cohort (www.ukbiobank.ac.uk) after phenotypic and genotypic quality control. The UKBB consortium initially recruited ~500 000 individuals aged 40–69 years (54% female) between 2006 and 2010. Phenotypic and cognitive measures were tested at the UKBB assessment centers or online, and also included demographic, socioeconomic and health data.

*Autism spectrum disorder cohorts*

We also included one cohort of children with autism spectrum disorder previously studied in Huguet et al., 2021[1]. In addition, we included 2,543 ASD probands with available IQ measures from the Simons Foundation Powering Autism Research (SPARK) database[2].

*Phenotypic measures*

We focus on the non-verbal IQ (NVIQ) when available, the fluid intelligence (FI) and on the g-factor otherwise. These intelligence measures were normalized using z-score transformations to render them comparable. We used the exact same process and data as shown previously in Huguet et al., 2021[1].

*Intelligence quotient*

In the SPARK cohorts, adapted tests have been used and ranked. We computed the average IQ interval for each rank to establish a numerical value. To be able to compare the different cognitive measures, all IQs were z-scored based on a mean of 100 and a standard deviation (SD) of 15.

*Fluid intelligence*

In UKBB, the FI score was assessed both in person (N=88,441, #20016) and online (N=13,773, #20191). This score is derived from 13 questions, measuring the capacity to solve problems requiring logic and reasoning abilities, independent of acquired knowledge. Participants were allotted 2 minutes to complete as many questions as possible from the test. The FI obtained were transformed into a z-score using the mean of 6.07 and the SD of 2.15 for the subgroup assessed in person, and using the mean of 6.61 and the SD of 1.98 for the subgroup assessed online.

*G-factor computation*

The g-factor is an indirect measure of general intelligence, obtained by extracting the first unrotated principal component from principal component analysis (PCA) of different standardized cognitive measures. It is a robust measure of general cognitive ability that is not very sensitive to the exact subtests used to calculate it as long as they measure a wide range of cognitive abilities[3]. Since cognitive measures used in the computation of the g-factor are not the same between tests used (in person and online), the g-factor was computed and normalized separately within each test group (in person and online) using the mean and SD computed on all available individuals. Of note, g-factor was computed before excluding individuals due to array quality control, leading to g-factors with means and SDs slightly different from 0 and 1 for the final subset of individuals included in our analyses.

For UKBB, the g-factor was computed using four cognitive tasks assessed in person (N=73,882) and online (N=62,080): Trail Making Test parts A and B (Executive function), Symbol Digit Substitution Test (Processing speed), Paired Associate Learning Test (Verbal declarative memory) and Picture Vocabulary (Crystallized ability) (Supplementary table 2). The g-factors obtained were transformed into a z-score using the mean of 1.80e-15 and the SD of 1.26 for the subgroup assessed in person, and using the mean of -1.40e-15 and the SD of 1.48 for the subgroup assessed online.

*Arrays processing and Genotyping*

UKBB used DNA extracted from blood and genotyped on two Affymetrix arrays (n=50k on UK BiLEVE and n=450k on UK Biobank Axiom)30 with ~95% probe overlap, using ~750k common markers. SPARK used DNA extracted from saliva (OGD-500 kit, DNA Genotek) genotyped on Illumina GSA-24v1-0 array (654k SNP sites).

*Genetic analysis on genotyping*

For data processing and quality control, we employed pLink[4] software. Each cohort was filtered to keep only autosomal SNPs with minor allele frequency (MAF) > 5%, probes providing genotypes that are not violating Hardy-Weinberg equilibrium (threshold $<1\times10\text{-}6$) and probes with call rates > 90%. Also, we used pLink[4] to check for duplicates, sex, and relationships for each participant with the same pipeline as previous work. Finally, we merged all genotyping data and ancestry (PC1 to PC10) was determined with KING[5] (with 3,615 common SNPs, we used the same quality control as in the previous step), we used the standard process defined in the website (https://www.kingrelatedness.com) and the referent population was 1000 Genomes.

*CNV calling*

We applied the same methodology as in Huguet et al. [1,6] available online (https://martineaujeanlouis.github.io/MIND-GENESPARALLELCNV/) on the array data using PennCNV[7] and QuantiSNP[8] algorithms. The following parameters were used for both algorithms: number of consecutive probes for CNV detection ≥3, CNV size ≥1Kb, confidence scores ≥15. CNVs detected by both algorithms were combined (CNVision) to minimize the number of potential false discoveries. After this merging step, an in-house algorithm based on CNV inheritance was applied to concatenate adjacent CNVs of the same type into one, according to the following criteria: a) gap between CNVs ≤150 kb; b) size of the CNVs ≥ 1000 bp; and c) number of probes ≥ 3.

*Array filtering*

After these steps, we remove from the analyses, all arrays for which a suspiciously high number of CNVs has been detected (≥ 50 for low resolution arrays [<1 million probes] and ≥ 200 for high resolution arrays [≥1 million probes]). For all cohorts, we used stringent quality-control criteria: call rate ≥95%; log R ratio-standard deviation <0.35; B allele frequency-standard deviation <0.08 and |wave factor|<0.05. The probes coordinates were updated from hg18 to hg19 using Illumina information and the liftover tool from the genome browser. From a total of 488,377 people with genotypic data, 28,522 were excluded for failing all these filters.

*Additional filtering to remove outliers*

All individuals with duplicated data or with discordant phenotypic and genetic information about the sex were removed (N=212). To avoid outliers which may add noise in our analyses, all carriers of a structural variant ≥ 10Mb, a mosaic CNV or with a chromosome anomaly (aneuploidy or sexual chromosome anomaly) were removed.

*CNV filtering*

After filtering the arrays according to their quality, we applied filtering for autosomal CNVs (X-linked CNVs were not investigated in this study). The CNVs with the following criteria were selected for analysis: confidence score ≥ 30 (for at least one of both detection algorithms), size ≥ 50 kb, unambiguous type (deletions or duplications) and overlap with segmental duplicates, HLA regions or centromeric regions < 50%, and at least ten probes for all array technologies used across the cohorts included in the analyses. Every recurrent CNV was manually visualized by at least one individual.

In addition, we applied an in-house algorithm based on a machine learning method to detect additional artifact CNVs. This algorithm was based on the consensus of three machine learning methods (Random forest, bagging of KNN and SVM) and on 9 CNV characteristics (Array criteria: log R ratio-standard deviation, B allele frequency-standard deviation, wave frequency; Localization CNV criteria: % of CNV overlap with centromeric regions and with segmental duplications; CNV criteria: density SNPs (size of CNV / numbers of SNPs), confidence score, % algorithms overlapping, type of CNV). This model was trained and tested respectively on 66% and 33% of a total of 34,156 CNVs (31,746 true CNVs and 2,410 artefacts from 6 cohorts, excluding SPARK), which represents a high confidence reference training set, and manually visualized by at least two individuals. Its application on the test set showed an AUC = 0.95, a sensitivity of 0.95 and a specificity of 0.85. This model was validated again on an additional naive dataset genotyped with another technology (GSA). We used 2,454 CNVs (1,936 true CNVs and 518 artefacts from SPARK cohort) and showed an AUC = 0.92, a sensitivity of 0.58 and a specificity of 0.97.

*Annotation of CNVs*

We annotated the CNVs using Gencode V19 annotation (hg19) with Ensembl gene name (*https://grch37.ensembl.org/index.html*). We used bedtools suite (*https://bedtools.readthedocs.io/en/latest/*) to identify the different elements of the genes encompassed in CNVs. Each coding gene with isoforms fully encompassed in filtered CNVs was annotated using the loss-of-function observed/expected upper bound fraction (LOEUF) score (gnomAD version 2.1.1)[9], which is available for 19,197 genes and ranges from 0.03 to 2. The smaller value defined gene intolerant to loss-of-function variants with cutoff on 0.35. CNV scores were derived by summing all scores (1/LOEUF) of genes within CNVs.

*Definition of gene-sets.*

Gene-sets are genes associated or preferentially expressed in various biological conditions. GOterm[10,11] from Ensembl (v.109, April 2023) and Synapse ontology from SynGO database[12] were used with propagated annotations following Gene Ontology Consortium recommendations. Genes overexpressed in adult[13,14] and fetal[15] brain cell types were fetched from Wagstyl et al[16]. Gene expressed in tissues, GTEx for whole body[17] and the Human Protein Atlas for brain (HPA)[18], were extracted from HPA version 22. Gene-sets by tissue are genes with an expression above 1SD compared to all tissue expressions. Throughout this study, we only considered gene sets containing more than 10 genes and disrupted by ≥ 30 CNVs covering at least 20% of the gene set.

*tagDS*

To test whether the deletion/duplication effect size ratio of a given gene-set deviates from the null model, we developed the trait-associated gene-dosage sensitivity score (tagDS). Each gene-set is represented in two dimensions by their deletion and duplication effect sizes. The nominal tagDS is the Euclidean distance between the gene-set coordinates and the line of equation, $y = 0.44x$ which is the ratio effect size between deletions (x) and duplications (y) computed for a genome-wide gene-set. Because effect-sizes are dependent of the gene-set sizes, we normalized this distance for each gene-set. Each nominal tagDS is then Z-scored based on the normal distribution of tagDS computing for 100 random gene-sets with the same number of genes.

*Linear regression model*

To estimate the effect of CNVs on cognitive ability, we used the model developed by Huguet at al.[1,6] based on linear regressions. The effect of genes completely encompassed in deletions or duplications on the Z-score of cognitive ability is computed by summing genes inside and outside gene-sets of interest (based on LOEUF, GOterm etc ...). Other covariates could be added depending on the model, nevertheless all models were adjusted by the 10 first ancestries PC. P-values were corrected for multiple testing (based on the biologic question and CNV type) using FDR correction. We fitted each model separately for deletion and duplication.

In our study, we utilized four distinct models to assess the average main effects of genes within specific categories of interest. In Model 1, the CNV-GWAS approach, we applied a linear model without adjusting for the multigenic burden in CNVs. Contrarily, in Models 2, 3, and 4, we also used linear models but incorporated adjustments for the effects of other genes within the CNV, accounting for variations in LOEUF values or gene-set outside the target window or gene-set.

**Model 1:** For each gene, we implemented an individual linear model, considering a minimum of 30 carriers. The aim was to assess the average main effect of each gene.

$$Z_{cog.\ ability\ adj.} \sim \beta_{0,DEL} + \beta_{1,\ DEL} \times Z_{gene\ was\ deleted\ 1\ or\ not\ 0.} + PC1\ to\ PC10$$

**Model 2:** Building on previously published work, we conducted 39 linear models to examine 38 overlapping LOEUF categories (using a sliding window with a size of 0.15 LOEUF), as well as a category comprising an ID gene list as defined by ClinGen (Supplementary Table 5). Each model focused on the average main effect of a gene within the specified category, adjusting for the impact of other genes in the CNV with LOEUF values falling outside the window of interest (Figure 2A).

$$Z_{cog.\ ability\ adj.} \sim \beta_0 + \beta_1 \times \sum(genes\ i\ inside\ the\ LOEUF\ window)$$

$$+ \beta_2 \times \sum(\ genes\ i\ with\ LOEUF < 1\ and\ outside\ the\ LOEUF\ window)$$

$$+ \beta_3 \times \sum(\ genes\ i\ with\ LOEUF \geq 1\ and\ outside\ the\ LOEUF\ window) + PC1\ to\ PC10$$

**Model 3:** We applied a linear model for each gene-set to estimate their effect sizes. These models evaluated the average main effects of genes within a gene-set, with adjustments for the influence of other genes in the CNV. Genes outside the gene-set, these were further subdivided into three categories: ID-gene, LOEUF < 1, and LOEUF ≥ 1.

$$Z_{cog.\,ability\,adj.} \sim \beta_0 + \beta_1 \times \sum(genes\ \ within\,the\,gene\,set)$$

$$+\ \beta_2\ \times \sum(ID\,genes\,i,\ outside\,the\,gene\,set)$$

$$+\ \beta_3 \times \sum(\,genes\,i\,with\,LOEUF\ <\ 1\,and\,outside\,the\,gene\,set)$$

$$+\ \beta_4 \times \sum(\,genes\,i\,with\,LOEUF \geq 1\,and\,outside\,the\,gene\,set) +\ PC1\ to\ PC10$$

**Model 4:** Employing the same approach as in Model 3, we used a single linear model but divided the gene-set into three LOEUF categories: LOEUF < 0.35, LOEUF in the range of [0.35, 1[, and LOEUF ≥ 1.

$$Z_{cog.\,ability\,adj.} \sim \beta_0 + \beta_1 \times \sum(\,genes\,i\,within\,the\,gene\,set\,and\,with\,LOEUF\ <\ 0.35)$$

$$+\ \beta_2 \times \sum(\,genes\,i\,within\,the\,gene\,set\,and\,with\,LOEUF\ =\ [0.35,\ 1[)$$

$$+\ \beta_3 \times \sum(\,genes\,i\,within\,the\,gene\,set\,with\,LOEUF \geq 1\,) +\ \beta_4\ \times \sum(ID\,genes\,i,\ outside\,the\,gene\,set)$$

$$+\ \beta_5 \times \sum(\,genes\,i\,with\,LOEUF\ <\ 1\,and\,outside\,the\,gene\,set)$$

$$+\ \beta_6 \times \sum(\,genes\,i\,with\,LOEUF \geq 1\,and\,outside\,the\,gene\,set) +\ PC1\ to\ PC10$$

### Supplementary tables

| Cohort | | Array type | N= | Ancestry | | Gender | | Age (year) | | Z-scored intelligence measure (adj) | | | Cognitive ability assessments |
|---|---|---|---|---|---|---|---|---|---|---|---|---|---|
| | | | | EUR | Others | F | M | Mean | SD | Mean | SD | Variables | |
| Unselected (n=258,292) | CaG | GSA | 2074 | 1982 | 92 | 1094 | 980 | 54.317 | 7.601 | 0.107 | 0.973 | Age, Age^2, sex, PC | g-factor, Reasoning, Memory, Reaction time |
| | | Omni2.5 | 515 | 490 | 25 | 281 | 234 | 52.437 | 8.602 | -0.009 | 0.956 | | |
| | | GSA + Omni2.5 | 2589 | 2472 | 117 | 1375 | 1214 | 53.943 | 7.845 | 0.084 | 0.970 | | |
| | G-Scot | 610Kq | 13715 | 13672 | 43 | 8081 | 5634 | 46.730 | 14.996 | 0.050 | 0.974 | Age, Age^2, sex, PC | g-factor, Logical Memory, Digit Symbol, Verbal fluency, Mill Hill Vocabulary |
| | Imagen | 610Kq; 660Wq | 1744 | 1624 | 120 | 891 | 853 | 14.450 | 0.366 | 0.441 | 0.977 | PC | WISC-IV (and g-factor, similarities score, vocabulary score, block design score, matrix reasoning score) |
| | LBC1936 | 610Kq | 503 | 500 | 3 | 246 | 257 | 69.825 | 0.829 | 0.047 | 0.946 | PC | Moray House Test |
| | SYS children | 610Kq | 559 | 557 | 2 | 298 | 261 | 15.058 | 1.894 | 0.361 | 0.882 | PC | WISC-III |
| | | HOE-12V | 408 | 408 | 0 | 207 | 201 | 14.906 | 1.760 | 0.212 | 0.848 | | |
| | | 610Kq + HOE-12V | 967 | 965 | 2 | 505 | 462 | 14.994 | 1.839 | 0.298 | 0.871 | | |
| | SYS parents | HOE-12V | 598 | 596 | 2 | 319 | 279 | 49.495 | 4.868 | -0.021 | 0.934 | Age, Age^2, sex, PC | g-factor, 12 cognitive measures‡ |
| | UKBB | Affymetrix | 73882 | 71364 | 2518 | 39317 | 34565 | 60.022 | 8.959 | 0.131 | 0.964 | Age, Age^2, sex, PC | g-factor [20] |
| | | | 62080 | 60484 | 1596 | 34335 | 27745 | 62.083 | 7.663 | 0.131 | 0.926 | | g-factor (online) |
| | | | 88441 | 80427 | 8014 | 47789 | 40652 | 58.139 | 8.304 | -0.035 | 0.961 | | FI |
| | | | 13773 | 13458 | 315 | 8284 | 5489 | 64.185 | 7.685 | -0.090 | 0.970 | | FI (online) |
| Autism (n=6,111) | SPARK | GSA | 2543 | 1984 | 559 | 540 | 2003 | 12.359 | 6.190 | -0.626 | 1.963 | PC | IQ |
| | MSSNG | WGS | 1007 | 768 | 239 | 202 | 805 | 9.503* | 4.600* | -0.529 | 1.590 | PC | IQ |
| | SSC | 1Mv1 | 332 | 279 | 53 | 44 | 288 | 9.538 | 3.240 | -0.602 | 1.558 | PC | WISC-IV n=19; DAS-II E-Y n=96; DAS-II S-A n=179; Mullen n=12; WASI-I n=26 |
| | | 1Mv3 | 1181 | 915 | 266 | 156 | 1025 | 8.769 | 3.523 | -0.982 | 1.638 | | WISC-IV n=16; DAS-II E-Y n=530; DAS-II S-A n=539; Mullen n=77; WASI-I n=19 |
| | | Omni2.5 | 1048 | 786 | 262 | 140 | 908 | 9.160 | 3.712 | -1.227 | 1.834 | | WISC-IV n=10; DAS-II E-Y n=403; DAS-II S-A n=494; Mullen n=124; WASI-I n=17 |
| | | 1Mv1 + 1Mv3 + Omni2.5 | 2561 | 1980 | 581 | 340 | 2221 | 9.028 | 3.576 | -1.033 | 1.722 | | WISC-IV n=45; DAS-II E-Y n=1,029; DAS-II S-A n=1,212; Mullen n=213; WASI-I n=62 |

**Supplementary table 1. Cohort descriptions:** Cohorts include 264,403 individuals, including 258,292 general populations. †63 and ‡ 12 cognitive measures were respectively used to compute the g-factor in SYS children and parents (Huguet et al 2021). SYS: Saguenay Youth Study, CaG: CARTaGEN, LBC1936: Lothian Birth Cohort 1936, SSC: Simons Simplex Collection; n=number of individuals remaining for analysis after quality control. The mean and Standard Deviation (SD) for g-factor slightly deviate from 0 and 1 in some cohorts since they were computed on all available data (before the exclusion of some individuals for poor quality array) and summarized here only for individuals included in the analyses. * The MSSNG cohort gave participants years but for not all, for 280 age was missing.

| field Name | Clinic value | | Online value | |
|---|---|---|---|---|
| | code | Lyall et al 2016[20] | code | Gfact 5 Online |
| Fluid intelligence score | 20016 | use | 20191 | use |
| Trail making #2 | 6350 | - | 20157 | use |
| Symbol digit substitution | 23324 | - | 20159 | use |
| Pairs matching | 399 | use | 20132 | use |
| Numeric memory | 4282 | use | 20240 | use |
| Prospective memory | 20018 | use | - | - |
| Mean time to correctly identify matches (Reaction time) | 20023 | use | - | - |

**Supplementary table 2. Descriptions of cognitive ability in UKBB were used.**

| Name | HPA | | | | GTEx | | | | Cell types | SynGO | LOEUF catg. | GO-term |
|---|---|---|---|---|---|---|---|---|---|---|---|---|
| | SD≥0.5 | SD≥1 | SD≥1.5 | SD≥2 | SD≥0.5 | SD≥1 | SD≥1.5 | SD≥2 | | | | |
| N lists | 215 | 215 | 215 | 215 | 37 | 37 | 37 | 37 | 29 | 85 | 38 | 6,130 |
| Mean list size | 3,817 | 1,975 | 1,017 | 571 | 3,056 | 2,015 | 1,173 | 787 | 890 | 120 | 1,423 | 89 |
| N unique coding gene | 12,710 | 12,706 | 12,449 | 12,427 | 12,755 | 12,758 | 12,751 | 12,538 | 8,422 | 921 | 13,288 | 11,460 |

**Supplementary table 3. Descriptions of Gene-set.**

**Supplementary table 4. CNV-GWAS details for each CNV (FDR) associated with cognitive ability.** (supplementary xls file)

**Supplementary Table 5: ClinGen 70 autosomal genes** (https://clinicalgenome.org/, supplementary xls file)

**Supplementary table 6: Summary statistics for meta-analyses.** (supplementary xls file)

**Supplementary table 7: Summary statistics for pooled-analyses.** (supplementary xls file)

**Supplementary table 8: SNPs observed with UCSC and GWAS-catalog inside chr2:109,510,927-110,376,563.** (supplementary xls file)

### Supplementary Figures

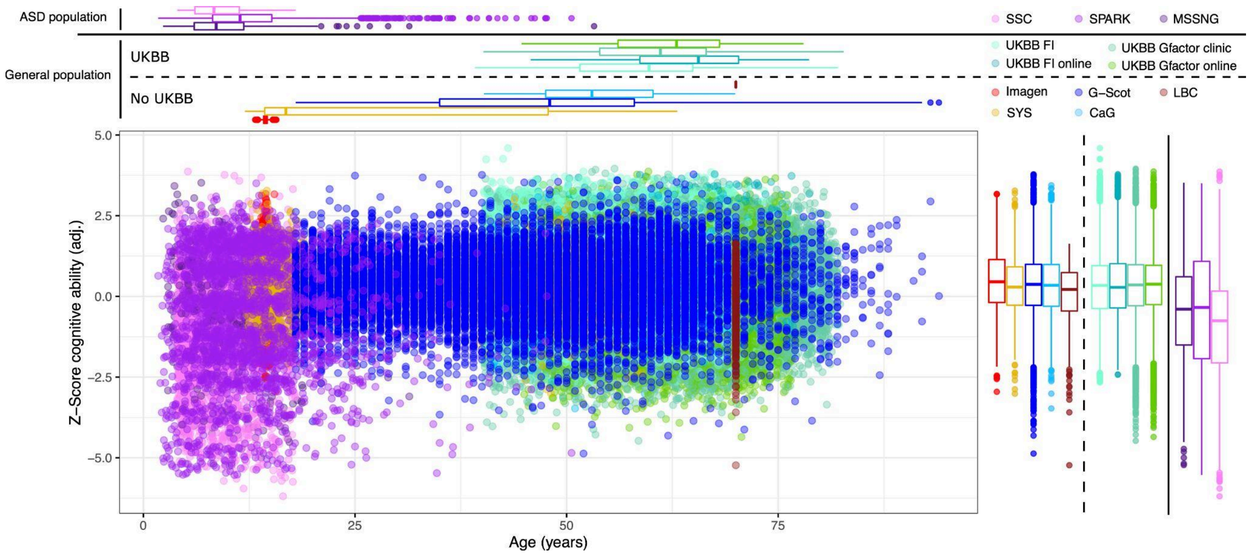


**Supplementary figure 1:** Description of cohorts in the dataset. The figure showed the cognitive ability z-score adjusted (sex, age, PC1 to 10 and cognitive test) with age for each participant.

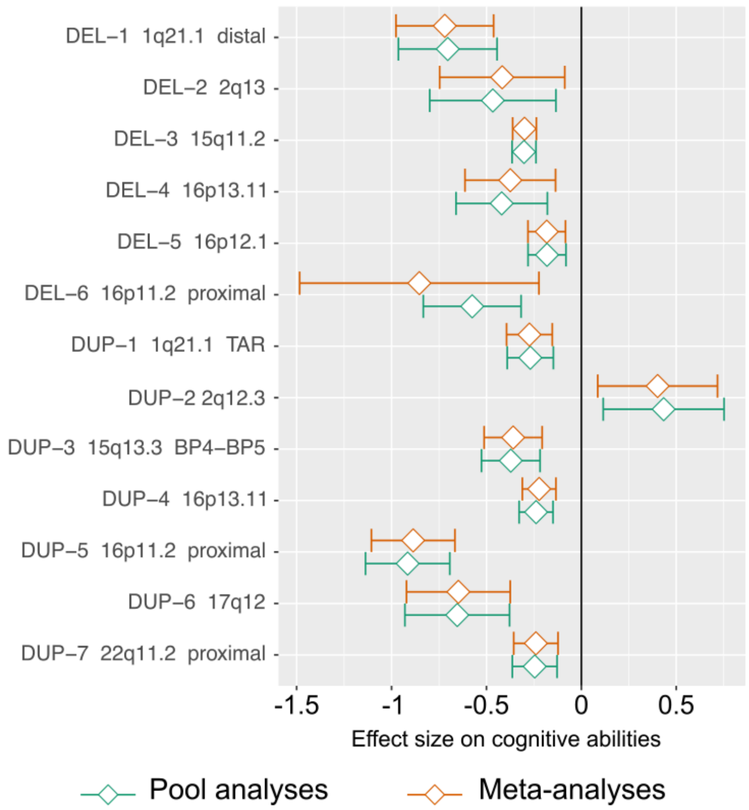


**Supplementary figure 2: Plot depicts the effect sizes for CNV-GWAS signals on cognitive abilities.** Green diamonds indicate pooled analyses (all cohorts regrouped), and orange diamonds represent meta-analyses (mean of effect sizes computed for each cohort separately). For meta-analyses, fixed-effect model values were chosen when the heterogeneity test was not significant ($p > 0.1$), and a random-effects model was employed when heterogeneity was significant. We displayed the values for the gene within the CNV that had the highest number of carriers.

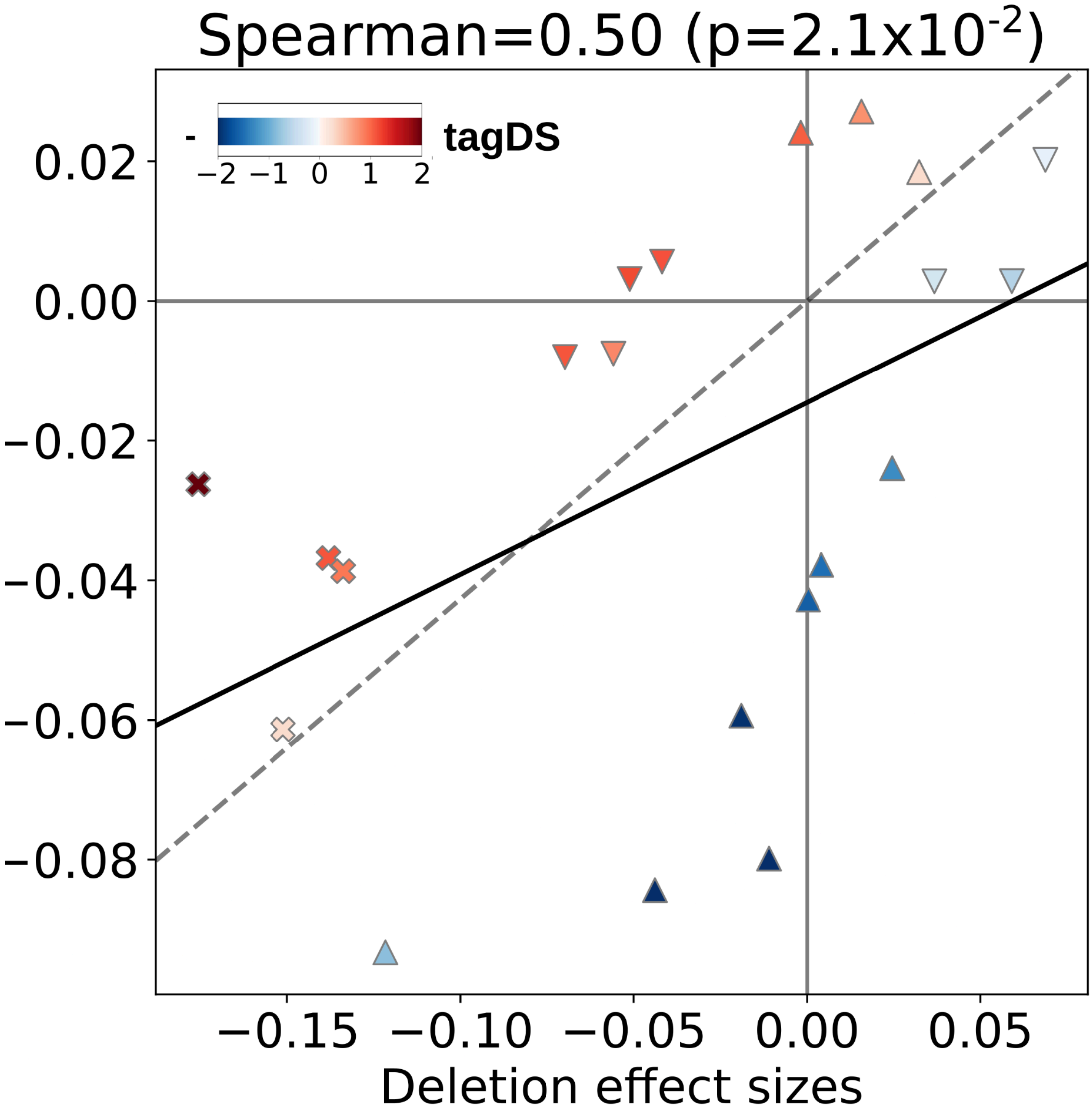


**Supplementary figure 3: Correlation on cognitive ability of LOEUF categories.** Spearman correlations (black line) between the effect sizes of deletions and duplications of gene-sets with FDR significant effects on cognitive ability for deletions (downward triangle), duplications (upward triangle), or both (cross). pvalue obtained from permutation test to account for the partial overlap between gene sets. Gene sets are color coded based on their tagDS. The dash line represents the average exome-wide duplication/deletion effect-sizes ratio.

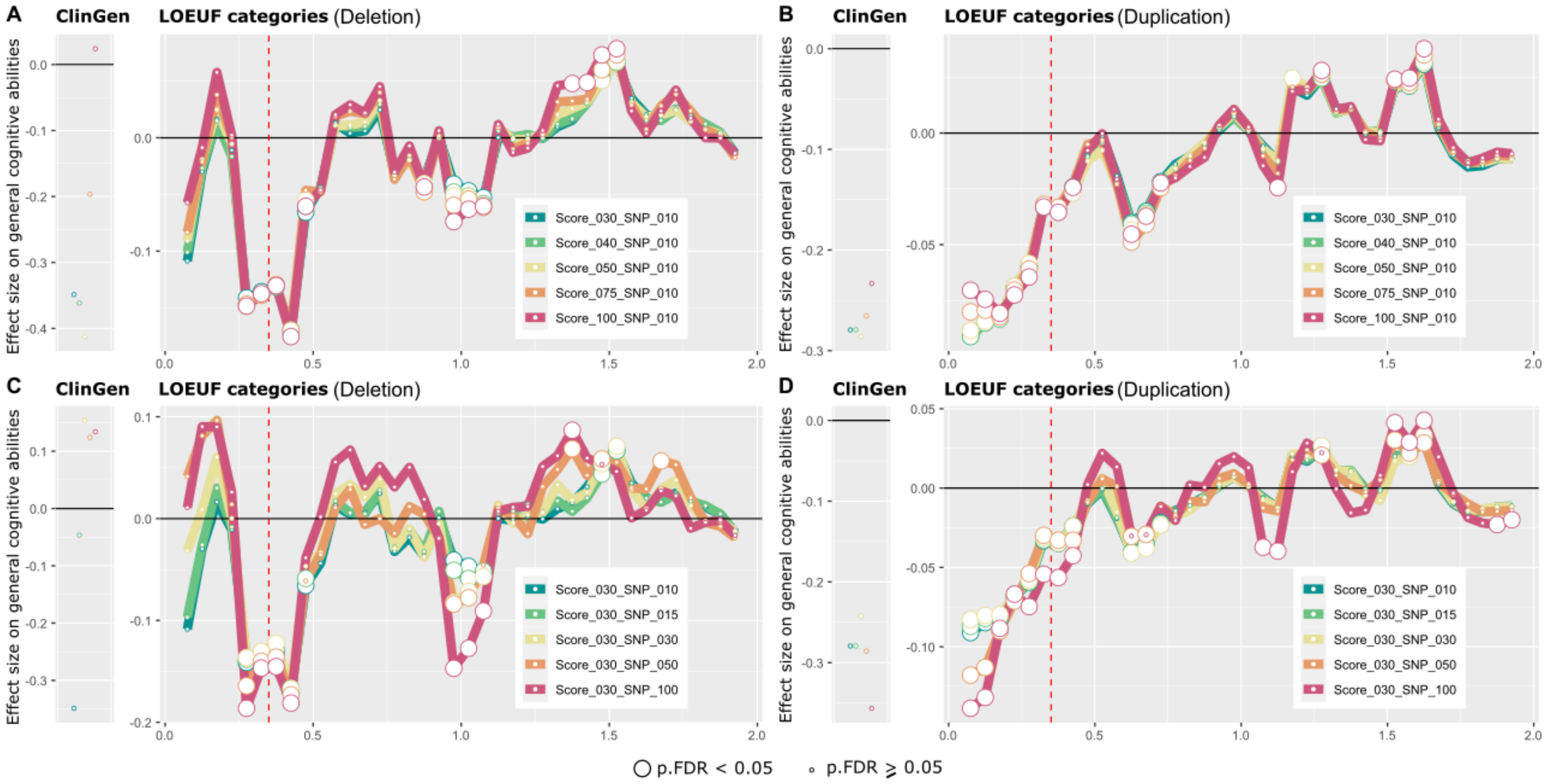


**Supplementary figure 4: Effect sizes of autosomal coding genes on general cognitive abilities based on their LOEUF values.** Sliding window estimating the effect size on cognitive ability of deletions (left) and duplications (right) for 38 LOEUF categories and definitive ID-genes curated by ClinGen (based on model 2). The line represents the estimated effect size of 38 categories of genes based on their LOEUF values in the model. Estimates were computed using a pooled dataset, large circles indicated significant pvalues adjusted by FDR and small one for non-significant. We ran sensitivity analyses based on different CNV cut-offs of quality controls with the likelihood score (Score ≥ 30, 40, 50, 75 and 100 and a fixed number of SNPs ≥10; for A and B) and numbers of SNPs inside CNV (SNPs ≥ 10, 15, 30, 50 and 100 and a fixed likelihood Score ≥30; C and D).

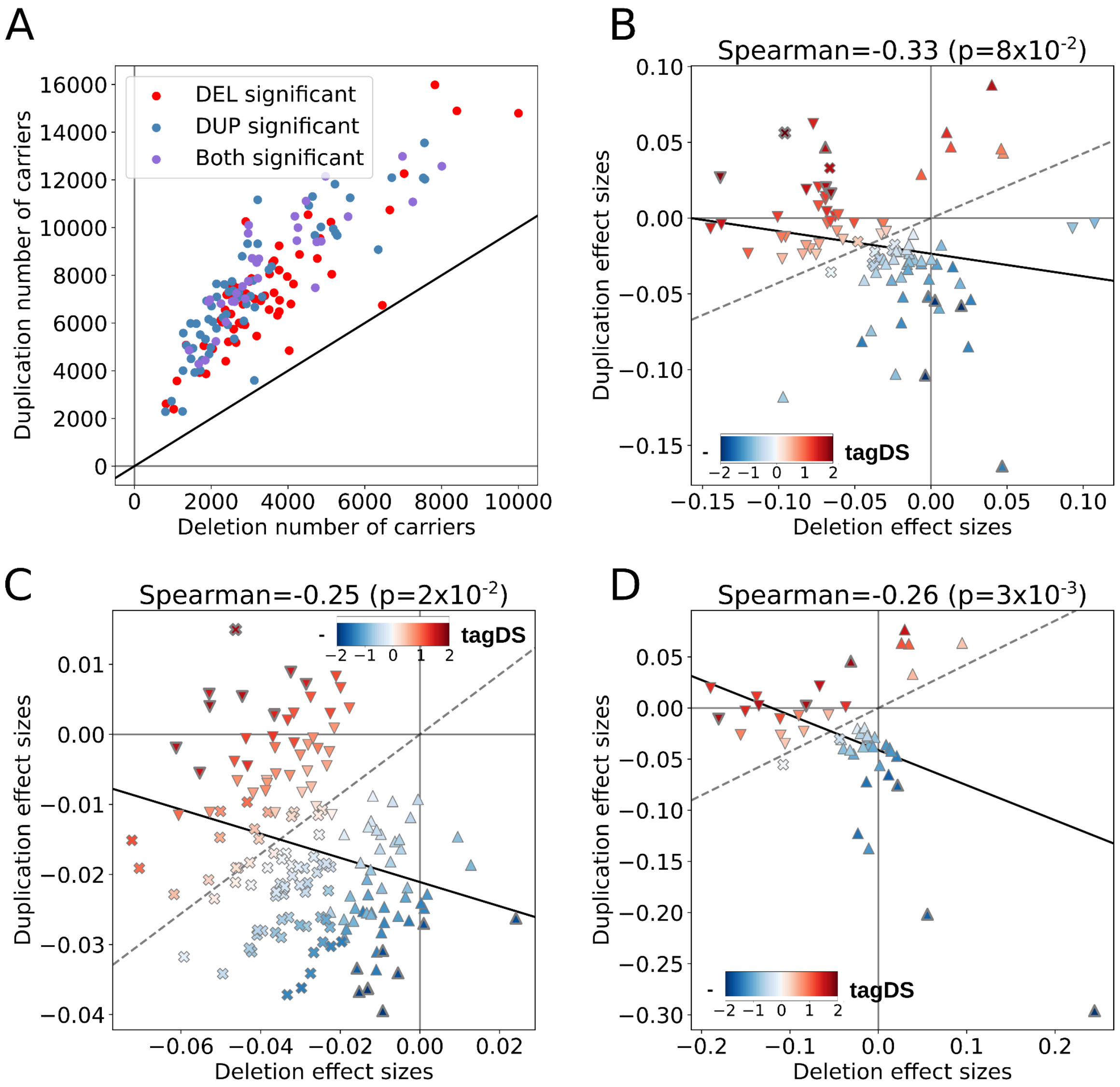


**Supplementary figure 5: Sensitivity analysis on cognitive ability for multiple HPA gene expression thresholds.** (A) Number of deletion and duplication carriers of genes for the 215 gene-sets analyzed in Figure 3. The black line represents the theoretical perfect concordance between Deletion and duplication carriers. Spearman correlations (black lines) between the effect sizes of deletions and duplications of tissue gene-sets with a normalized expression threshold >0.5SD (B), >1.5SD (C), and >2SD(D). FDR significant effects on cognitive ability for deletions (downward triangle), duplications (upward triangle), or both (cross). pvalues obtained from permutation tests to account for the partial overlap between gene sets. Gene sets are color coded based on their tagDS. The dash line represents the average exome-wide duplication/deletion effect-sizes ratio.

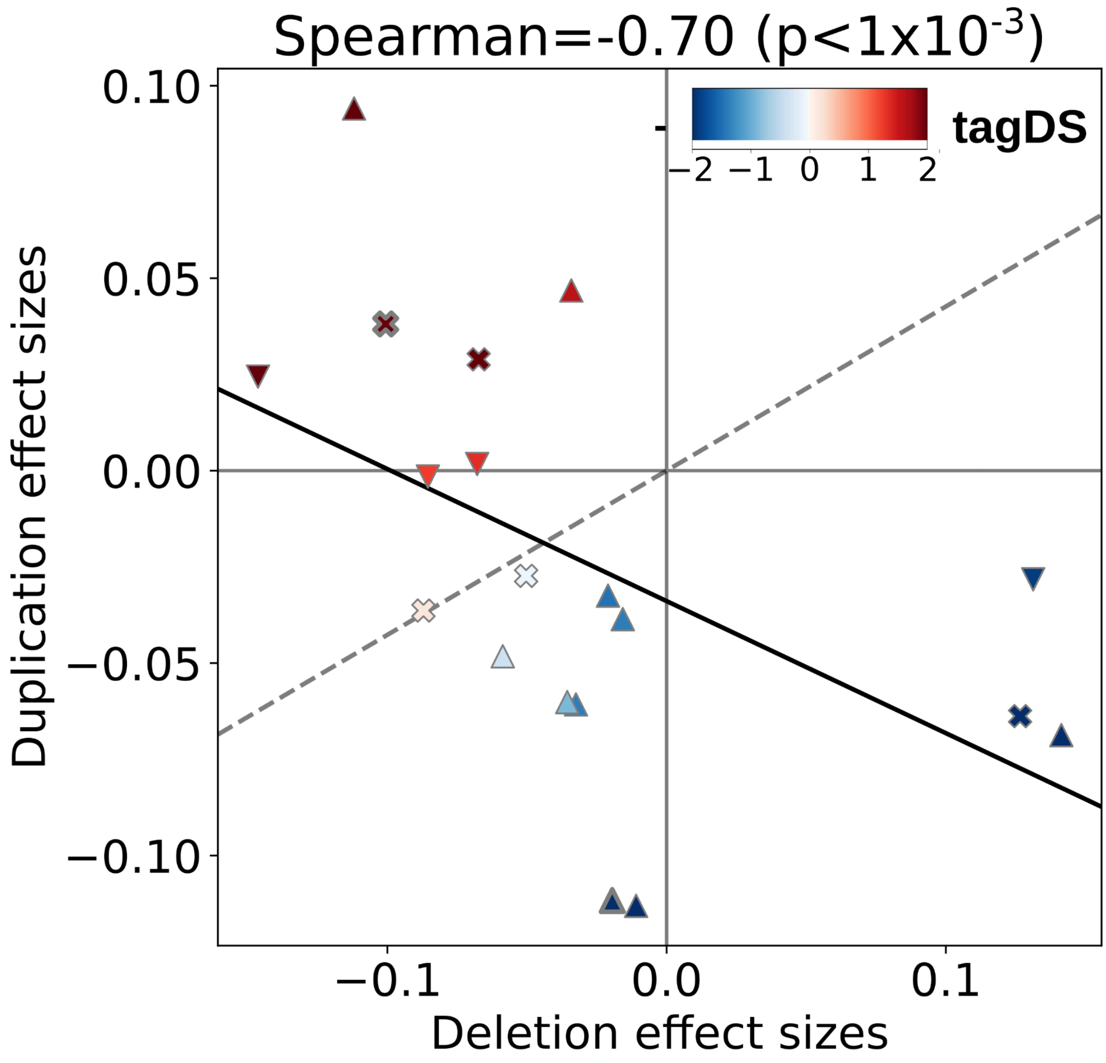


**Supplementary figure 6: Correlation on cognitive ability of cell type.** Spearman correlation (black line) between the effect sizes of deletions and duplications of gene-sets with FDR significant effects on cognitive ability for deletions (downward triangle), duplications (upward triangle), or both (cross). pvalues obtained from permutation tests to account for the partial overlap between gene sets. Gene sets are color coded based on their tagDS. The dash line represents the average exome-wide duplication/deletion effect-sizes ratio.

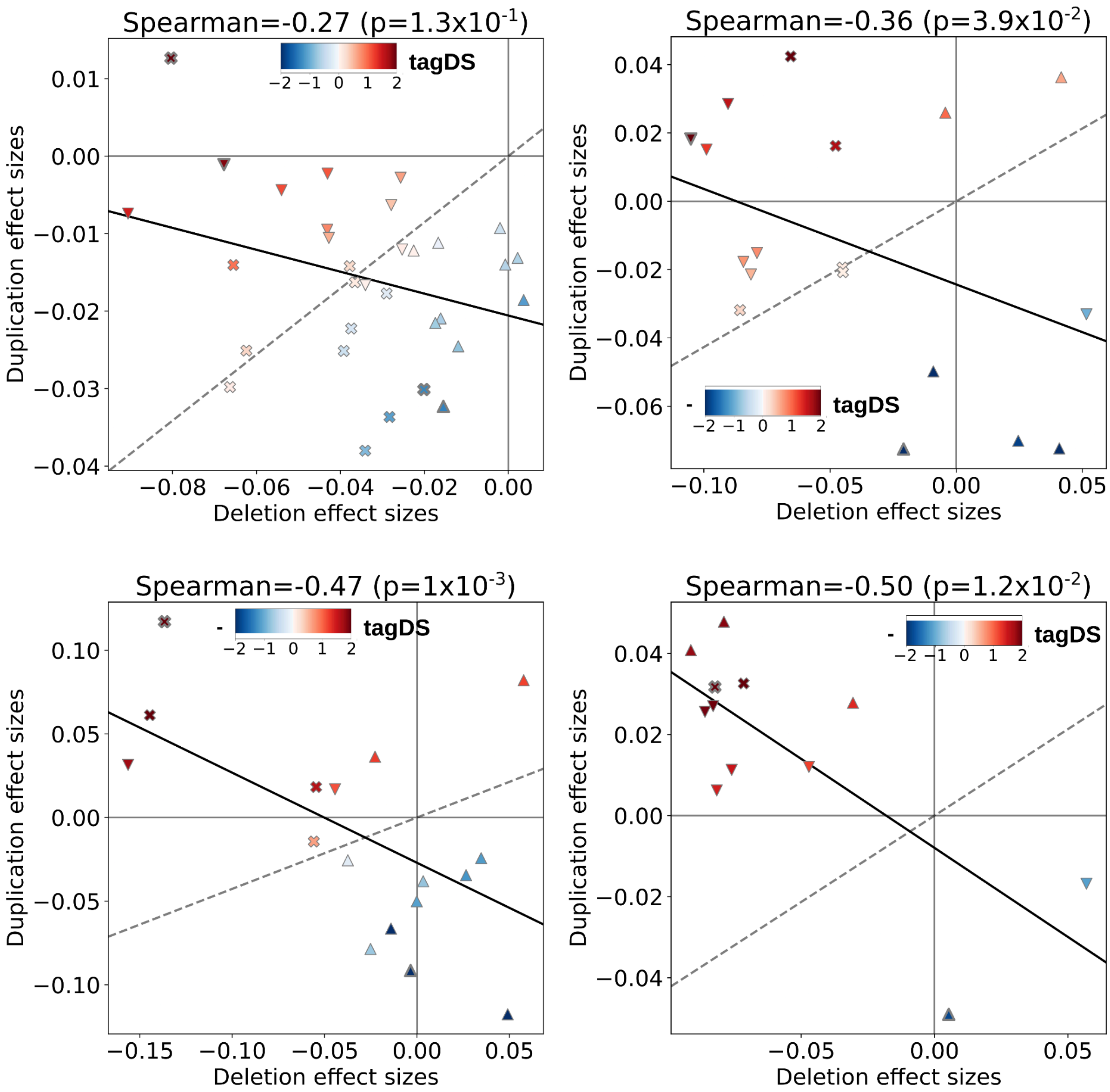


**Supplementary figure 7: Correlations on cognitive ability of multiple GTEx gene specificity thresholds.** Spearman correlations (black lines) between the effect sizes of deletions and duplications of tissue gene-sets with a normalized relative expression threshold >0.5SD (A), >1.5SD (B), >2SD(C) and >1SD without low-tissue-specificity genes(D). FDR significant effects on cognitive ability for deletions (downward triangle), duplications (upward triangle), or both (cross). pvalues obtained from permutation tests to account for the partial overlap between gene sets. Gene sets are color coded based on their tagDS. The dash line represents the average exome-wide duplication/deletion effect-sizes ratio.

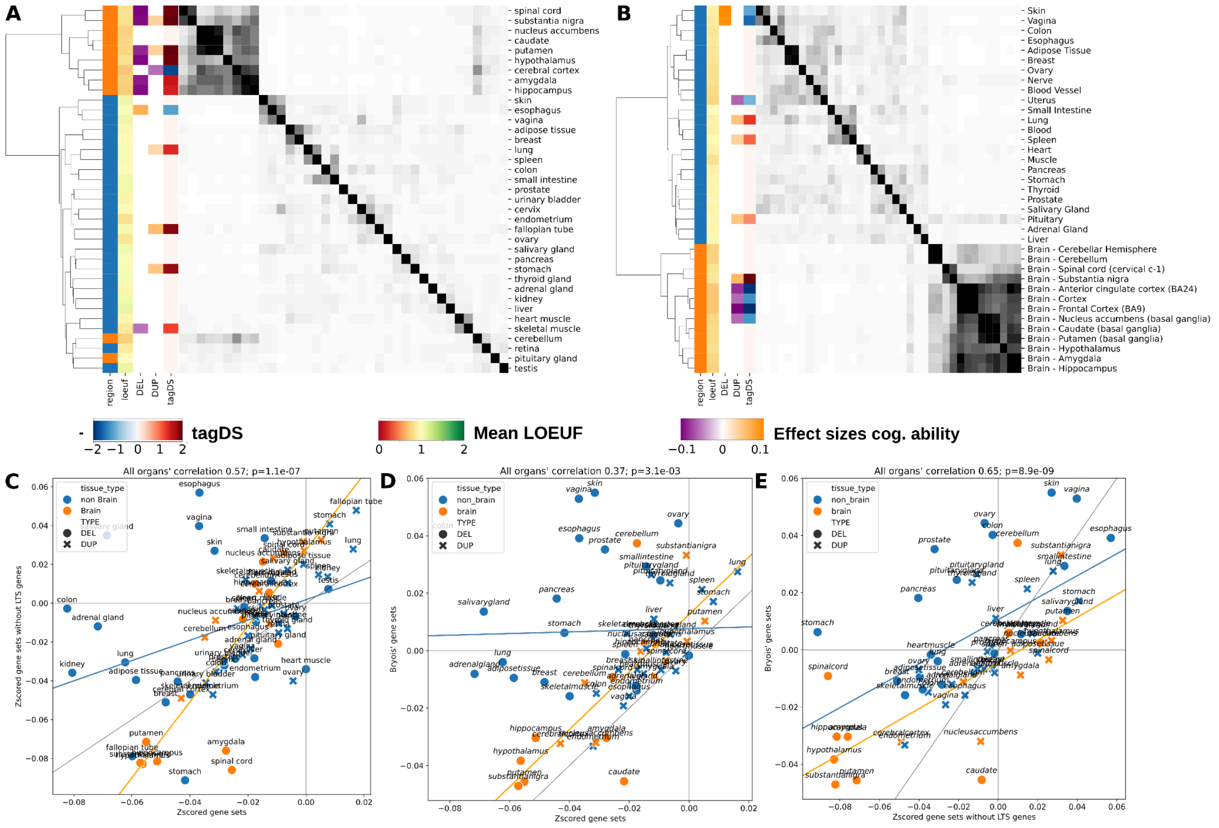


**Supplementary figure 8: Comparisons of effect sizes for different gene-set associations.** Clustering of gene-sets computed with z-score minus 8,194 low-tissue-specificity genes (A) and computed with TDEP (Bryois et al) (B). Orange represents brain tissues and blue non-brain tissues. Gene-set overlap matrix show high overlap between brain gene-sets and much lower overlap across non-brain tissues for both association methods. 2nd columns represent the average LOEUF score of the gene-set. 3rd and 4th columns represent the effect of gene-sets on cognitive ability when deleted and duplicated, respectively. The 5th column is the resulting tagDS. Correlation of effect-sizes for multiple gene-set definitions, z-scored gene-sets with all coding genes versus z-scored gene-sets without low-tissue-specificity genes (C), z-scored gene-sets with all coding genes versus TDEP gene-sets (D) and z-scored gene-sets without low-tissue-specificity genes versus TDEP gene-sets (E). Orange lines show the Spearman correlation between brain gene-sets (r=0.84 p=1e-6; r=0.82 p=9e-6; r=0.76 p=1e-4 for B, C and D respectively) and blue lines show correlation for non-brain gene-sets (r=0.47 p=4e-4; r=0.16 p=3e-1; r=0.58 p=4e-5 for B, C and D respectively). The grey line represents a theoretical perfect concordance between estimates. Pvalues are nominal pvalues without permutation tests.

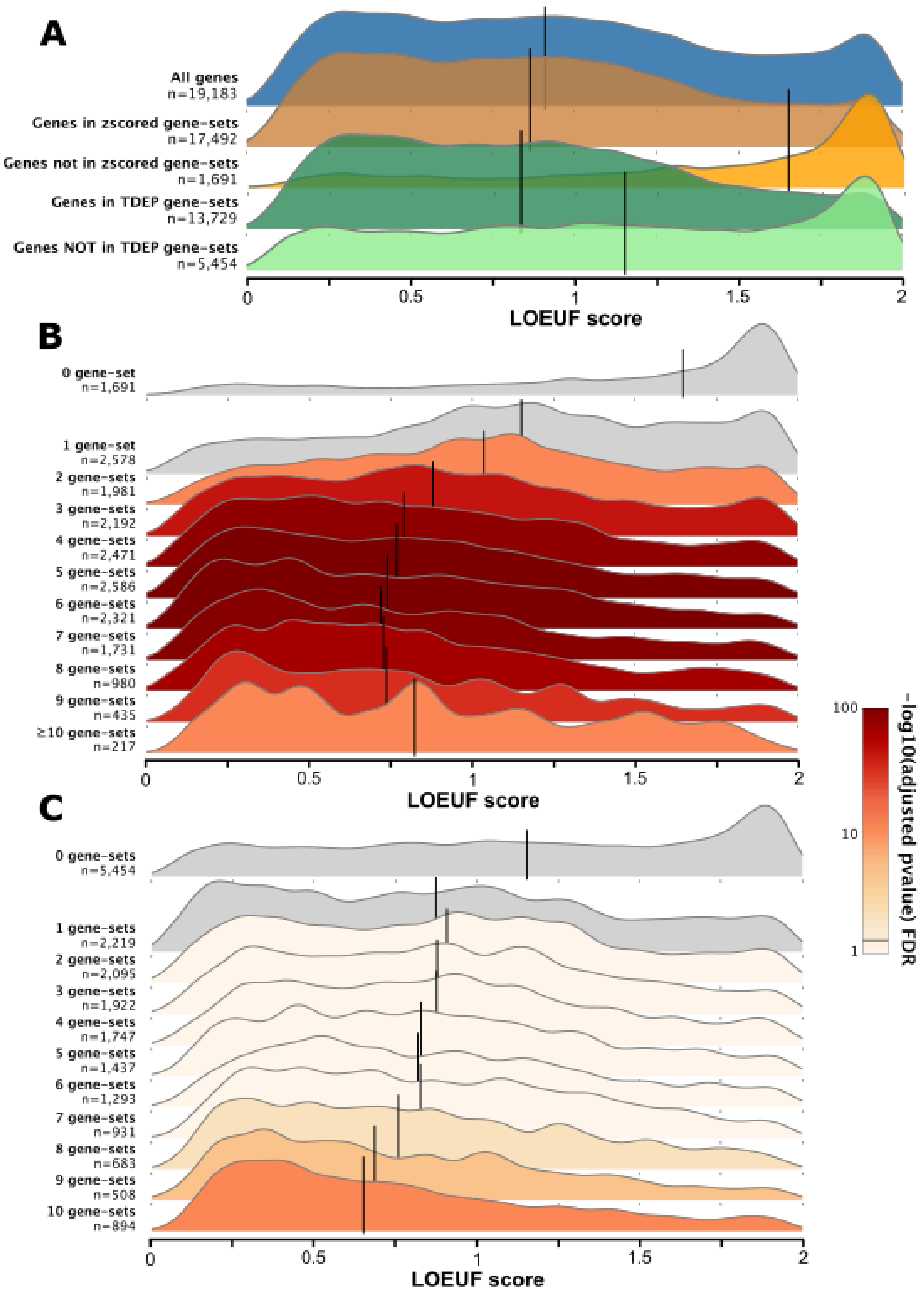


**Supplementary figure 9: Ridgeplots representing the LOEUF distribution across multiple levels of specificity.** (A) Distribution of LOEUF values for the whole coding exome, for genes assigned to at least one GTEx gene-set defined by z-score, for genes not assigned to any GTEx tissue (Z-score), for genes assigned to at least one GTEx tissue defined by TDEP and for genes not assigned to any GTEx gene-set (TDEP). Distribution of LOEUF for genes present in one or multiple gene-sets defined by Z-score (B) and TDEP (C) methods. The color represents the FDR-adjusted pvalue of the Mann-Whitney test between the distribution of LOEUF for specific genes (present in only one gene-set) and the distribution of interest. Black line represents the median LOEUF score for each category.

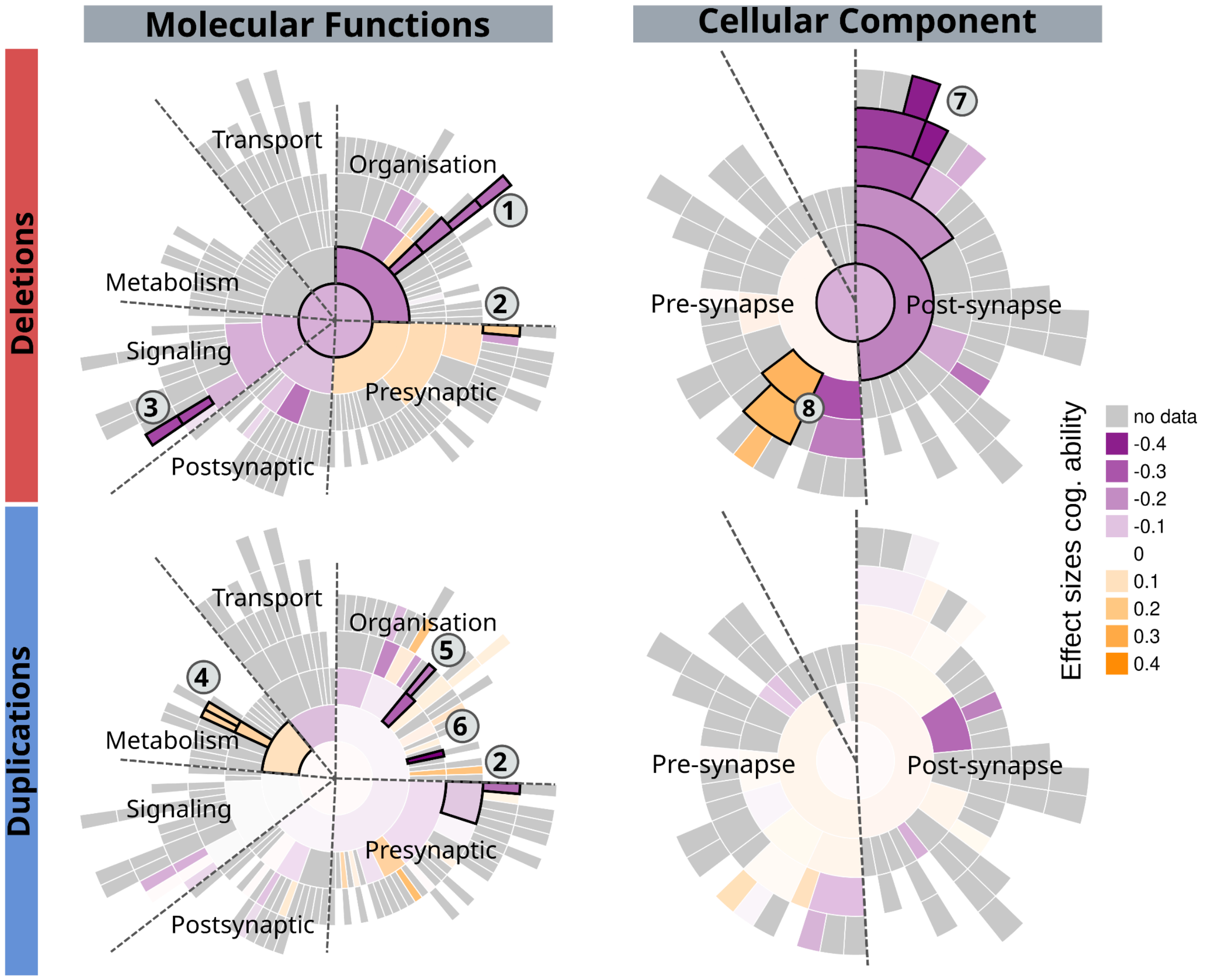


**Supplementary figure 10: Effect sizes of Deletion and duplication on cognitive ability for SynGO gene-sets.** Effect sizes of synaptic molecular functions and cellular component gene-sets as defined by SynGO[12] on cognitive ability. Purple and orange represent negative and positive effect size on cognitive ability, respectively. Ontologies with black edges indicate significant effects (FDR). The results are shown only for SynGO terms with more than 10 genes, observed at least 30 times in our dataset, and with a coverage greater than 20%. Note: 1) Regulation of modification of postsynaptic actin cytoskeleton, 2) Regulation of calcium-dependent activation of synaptic vesicle fusion, 3) Presynaptic modulation of chemical synaptic transmission, 4) translation at synapse, 5) regulation of postsynapse organization, 6) synapse adhesion between pre- and post-synapse, 7) Integral component of postsynaptic density membrane, 8) Synaptic vesicle membrane.

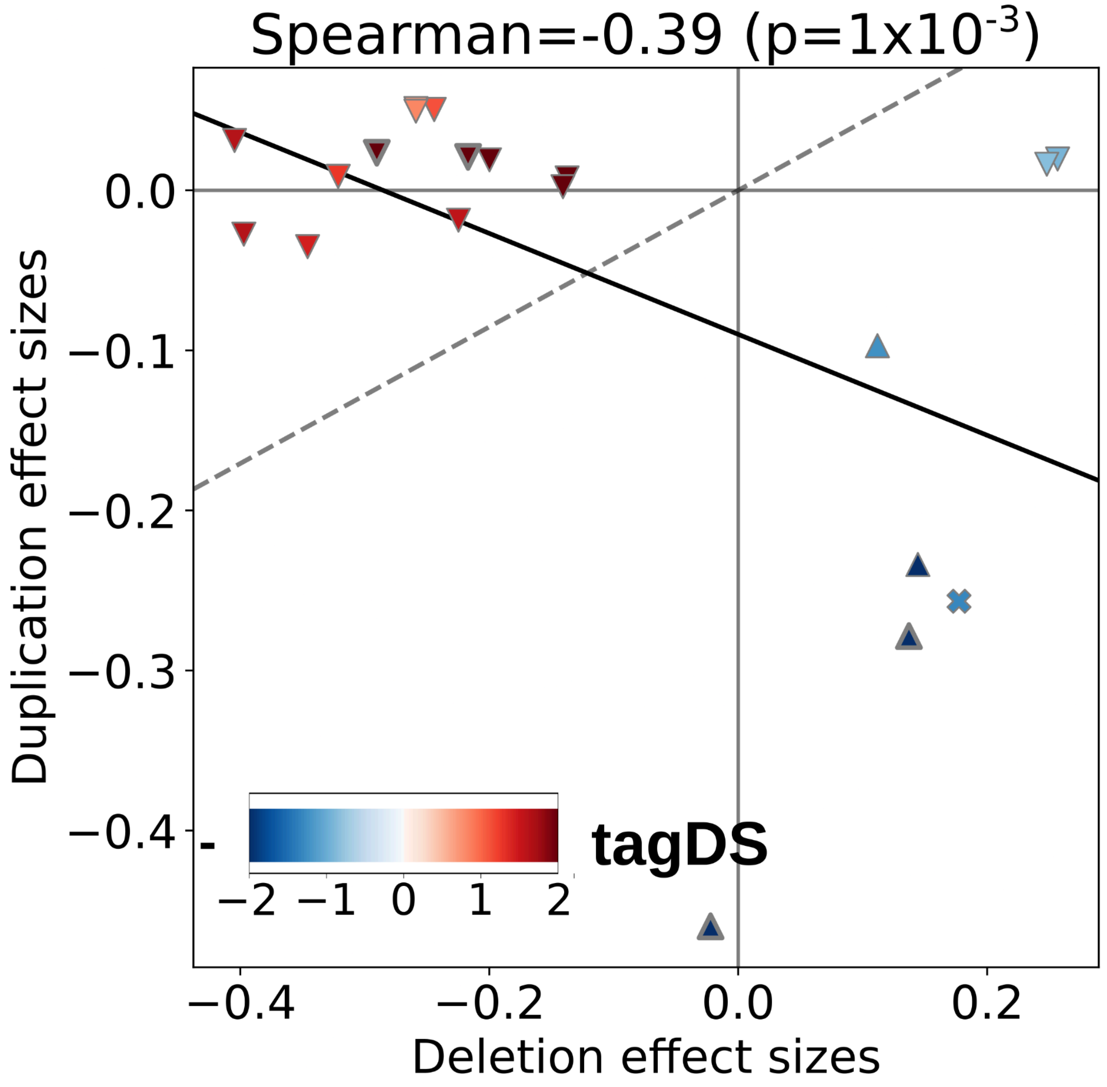


**Supplementary figure 11: Correlation on cognitive ability of SynGO.** Spearman correlations between the effect sizes of deletions and duplications of gene-sets with FDR significant effects on cognitive ability for deletions (downward triangle), duplications (upward triangle), or both (cross). pvalues obtained from permutations to account for the partial overlap between gene sets. Gene sets are color coded based on their tagDS.

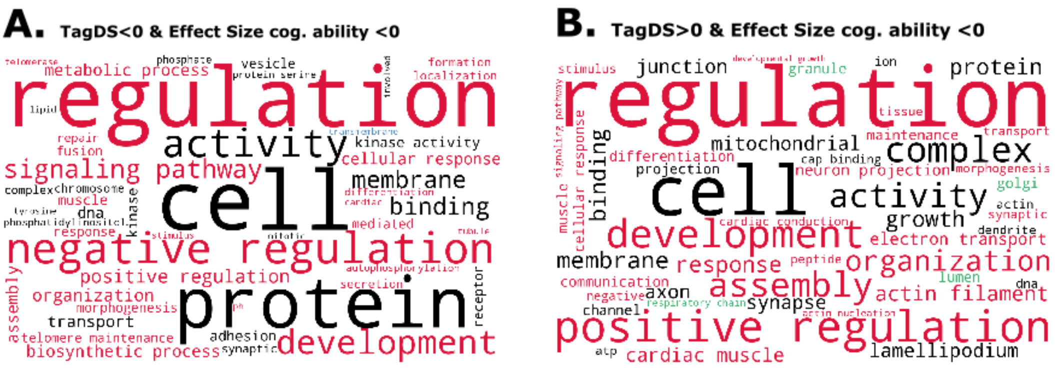


**Supplementary figure 12: Word-cloud plots on effects on cognitive ability of gene clustered, based on Gene ontology annotation.** Word-cloud plots (A) and (B) display the 50 most frequent words within GO-term names and having a negative effect on cognitive abilities, with negative (n GO-term = 279) and positive ($n_{GO\text{-}term}$ = 242) tagDS, respectively. Red, blue, and green words represent biological processes, molecular functions and cellular components terms, respectively. Black words are observed in multiple categories.

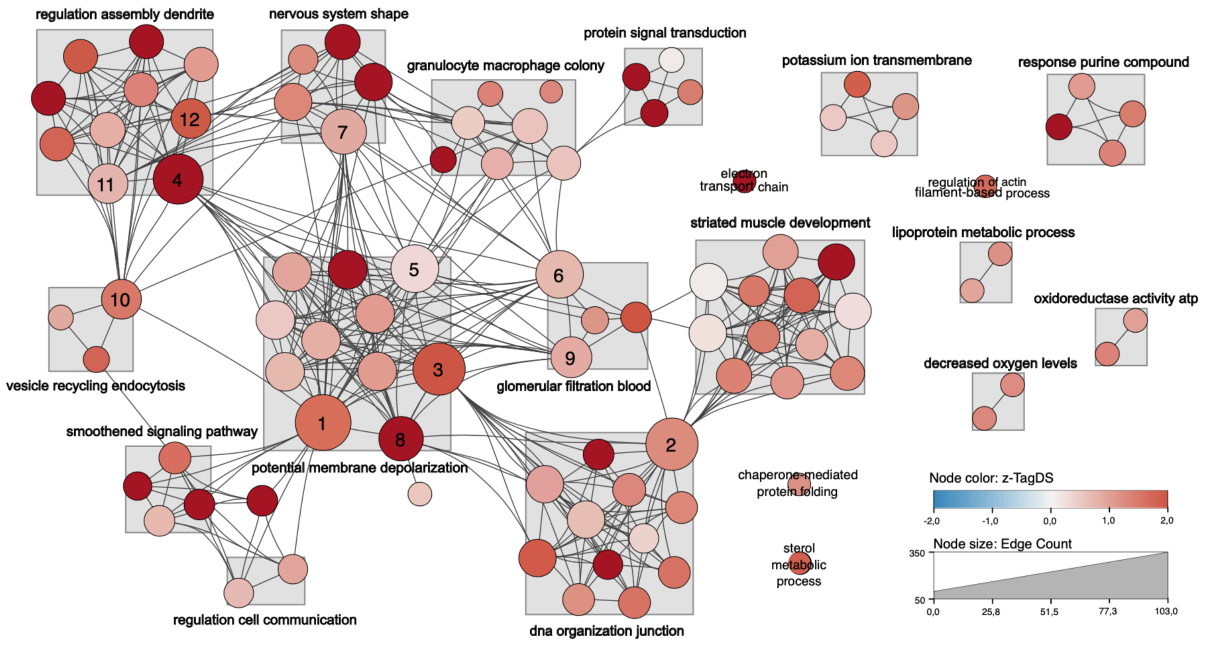


**Supplementary figure 13: Network of GO-Terms associated with positive tagDS and negative impact on cognitive ability.** This figure presents a Revigo-generated[21] network of GOterms based on GOterm associted with negative impact on cognitive ability and positive tagDS. Each node symbolizes a specific GOterm or GOterm cluster (clustering by standard Revigo criteria[21]). Node color denotes the tagDS value associated (red for positive tagDS). The node size of each node correlates with its edge count, reflecting the extent of its connectivity and relevance within the network. Larger nodes represent higher numbers of interactions with other GO-terms. Links between nodes depict the connection between GO-Terms. The bold text represents the supra-cluster defined by ReviGO[21], represented by a grey square. Nodes had at least 15 edge counts: 1) positive regulation of excitatory postsynaptic potential; 2) axonogenesis; 3) regulation of cellular component size; 4) regulation of synapse organization; 5) positive regulation of vasoconstriction; 6) regulation of glomerular filtration; 7) regulation of cell shape; 8) cell volume homeostasis; 9) negative regulation of blood pressure; 10) positive regulation of endocytosis; 11) positive regulation of dendrite development; 12) regulation of dendrite development.

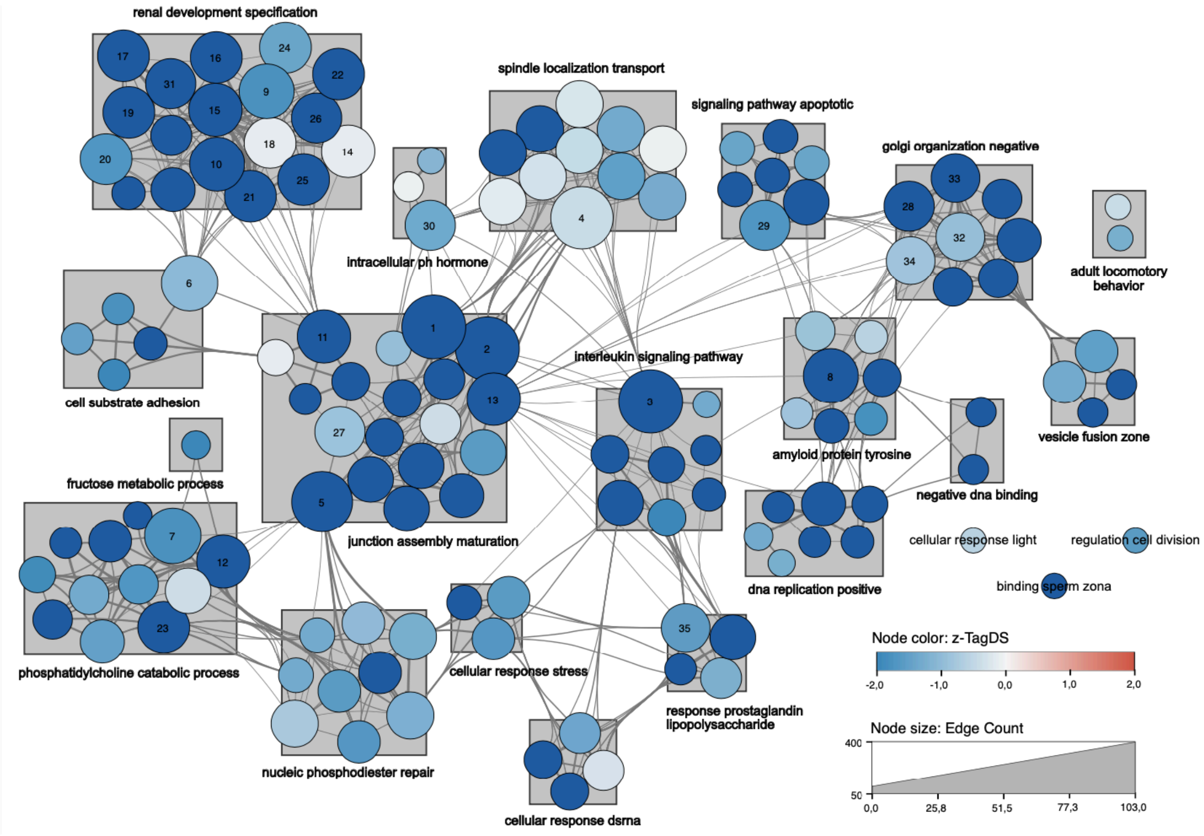


**Supplementary figure 14: Network of GO-Terms associated with negative tagDS and negative impact on cognitive ability.** This figure presents a Revigo-generated[21] network of GOterms based on GOterm associted with negative impact on cognitive ability and negative tagDS. Each node symbolizes a specific GOterm or GOterm cluster (clustering by standard Revigo criteria[21]). Node color denotes the tagDS value associated (blue for negative tagDS). The node size of each node correlates with its edge count, reflecting the extent of its connectivity and relevance within the network. Larger nodes represent higher numbers of interactions with other GO-terms. Links between nodes depict the connection between GO-Terms. The bold text represents the supra-cluster defined by ReviGO[21], represented by a grey square. Nodes had at least 15 edge counts: 1) vesicle fusion; 2) phagolysosome assembly; 3) signal release; 4) establishment of spindle localization; 5) telomere maintenance; 6) neuron migration; 7) NAD metabolic process; 8) negative regulation of translation; 9) cardiocyte differentiation; 10) cardiac neural crest cell development involved in heart development; 11) synapse maturation; 12) nucleoside phosphate biosynthetic process; 13) mitotic G2/M transition checkpoint; 14) muscle tissue development; 15) formation of primary germ layer; 16) trachea development; 17) somite development; 18) skeletal muscle organ development; 19) segmentation; 20) renal tubule development; 21) embryonic pattern specification; 22) cerebellar cortex development; 23) ceramide biosynthetic process; 24) blastocyst formation; 25) axis specification; 26) trachea morphogenesis; 27) nuclear division; 28) negative regulation of cell projection organization; 29) negative regulation of apoptotic signaling pathway; 30) hormone transport; 31) anterior/posterior axis specification; 32) negative regulation of supramolecular fiber organization; 33) negative regulation of neuron projection development; 34) negative regulation of cytoskeleton organization; 35) lipopolysaccharide-mediated signaling pathway.

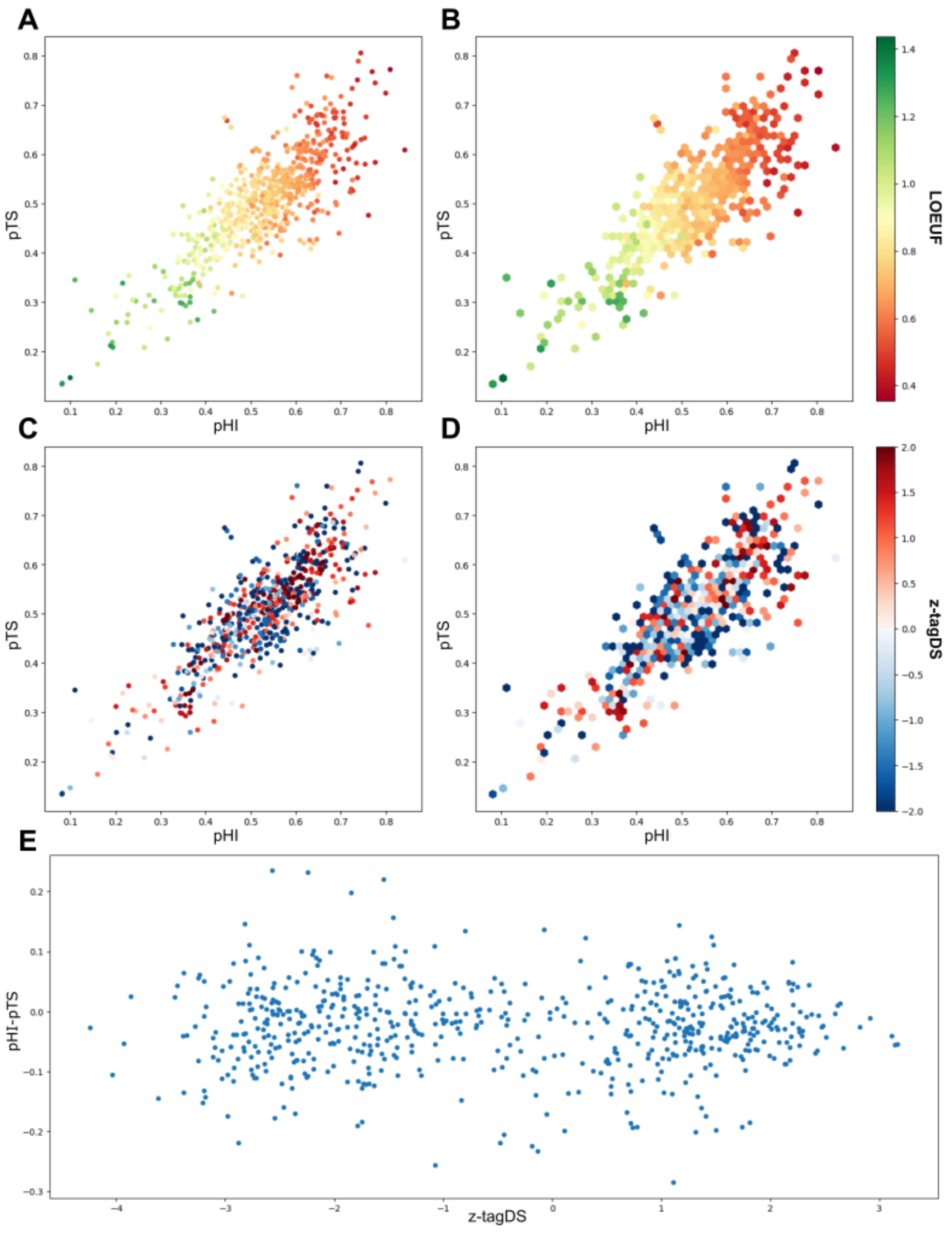


**Supplementary figure 15: Comparison between Collins et al.'s hapoinsufficiency, triplosensitive scores and tagDS.** Scatter (A) and hexbin (B) plots of the distribution of mean pHI and pTS of genes coming from the 645 significant GO-terms. The color-code represent the average LOEUF value of GO-terms. Hexbin plots visualize the average between overlapping points. Scatter (C) and hexbin (D) plots representing again the distribution of mean pHI and pTS of genes coming from the 645 significant GO-terms. The color-code represent the z-tagDS value extract from Figure 6. (E) Comparison between gene dosage specificity for cognitive ability as describe by z-tagDS and gene dosage specificity based on pHI, pTS difference.
